# Components of micro-evolutionary and phenotypic change in seasonal migration versus residence in a wild population

**DOI:** 10.1101/2022.12.31.522097

**Authors:** Paul Acker, Francis Daunt, Sarah Wanless, Sarah J. Burthe, Mark A. Newell, Michael P. Harris, Robert L. Swann, Carrie Gunn, Tim I. Morley, Jane M. Reid

## Abstract

Dissecting joint micro-evolutionary and plastic responses to environmental perturbations fundamentally requires quantifying interacting components of genetic and environmental variation underlying expression of key traits. This ambition is particularly challenging for phenotypically discrete traits where multiscale decompositions are required to handle non-linear transformations of underlying genetic and environmental variation into phenotypic variation, especially when effects have to be estimated from incomplete field observations. We devised a novel joint multistate capture-recapture and quantitative genetic animal model, and fitted this model to full-annual-cycle resighting data from partially migratory European shags (*Gulosus aristotelis*) to estimate key components of genetic, environmental and phenotypic variance in the ecologically critical discrete trait of seasonal migration versus residence. We demonstrate non-trivial additive genetic variance in latent liability for migration, resulting in estimated micro-evolutionary responses following two episodes of strong survival selection. Yet, underlying additive genetic effects interacted with substantial permanent individual and temporary environmental effects to generate complex non-additive effects, causing large intrinsic gene-by-environment interaction variance in phenotypic expression. Our findings reveal how temporal dynamics of seasonal migration result from combinations of instantaneous micro-evolution and within-individual phenotypic inertia, and highlight how plastic phenotypic variation could expose cryptic genetic variation underlying discrete traits to complex forms of selection.

## INTRODUCTION

Understanding short-term and longer-term population responses to environmental perturbations and changes, manifested through combinations of micro-evolution and plasticity, requires quantifying multiple components of variance underlying key traits and disentangling their independent and interacting effects (Sgrò et al. 2016; Bay et al. 2017; Kingsolver and Buckley 2017; Kelly 2019; Hansen and Pélabon 2021). Specifically, the additive genetic variance defines the potential for micro-evolutionary responses to selection and hence for inherited cross-generational changes in phenotypic distributions. Further, together with any permanent individual variance stemming from lifelong effects of developmental environments and/or non-additive genetic effects, the additive genetic variance can also shape the degree of short-term phenotypic inertia due to repeatability, thereby generating potential for within-generation changes resulting from survival selection. Meanwhile, variance attributable to temporary effects of current environments encompasses rapid phenotypic responses through labile plasticity. Quantifying these components, and pinpointing their joint effects on phenotypic dynamics, can therefore reveal how population responses to environmental change are shaped over multiple timeframes (Gienapp et al. 2008; Merilä and Hendry 2014; Pigeon et al. 2017; Bonnet et al. 2019). However, major conceptual and analytical advances are still required to quantify and interpret complex effects and interactions that can generate non-linear constraints on phenotypic expression and micro-evolutionary dynamics in wild populations (Morrissey 2015; Hansen and Pélabon 2021; Chevin et al. 2022).

Such challenges particularly apply for key life-history traits that are expressed as discrete alternative states rather than as continuously distributed phenotypes (e.g. movement vs. philopatry; breeding vs. non-breeding; dormancy vs. activity; survival vs. death; Caswell 2001; Reid and Acker 2022). Such traits are commonly shaped by numerous genetic and environmental effects, and hence are appropriately treated within the broad framework of quantitative genetics. An explicit model is therefore needed for how linear combinations of effects generate non-linear (dichotomous) phenotypic outcomes (i.e. non-linear genotype-environment-phenotype map; Houle et al. 2010; Chevin et al. 2022; Brun-Usan et al. 2022). This is parsimoniously achieved by the threshold trait model, which is empirically based and has long been successfully utilised to selectively breed discrete traits in agricultural and laboratory systems (Wright 1934; Gianola 1982; Roff 1996; Lynch and Walsh 1998; Moorad and Promislow 2011). Here, dichotomous alternative phenotypes are expressed when an underlying latent continuously distributed liability is above versus below some threshold (Figure 1a). Accordingly, partitioning variance into summed additive genetic, permanent individual and temporary residual effects can be appropriately achieved on the liability scale (Falconer and McKay 1996; de Villemereuil et al. 2016). However, emerging phenotypic variation, on which selection will act, is structurally different because phenotypic expression is bounded by the discrete possible outcomes. Strictly additive effects on liabilities can therefore generate non-additive effects on phenotypes. Further, such non-additive effects increase with distance of underlying liabilities from the threshold, causing intrinsic links between the liability mean and the phenotypic variance (Figure 1; Dempster and Lerner 1950; Reid and Acker 2022).

**Figure 1.**
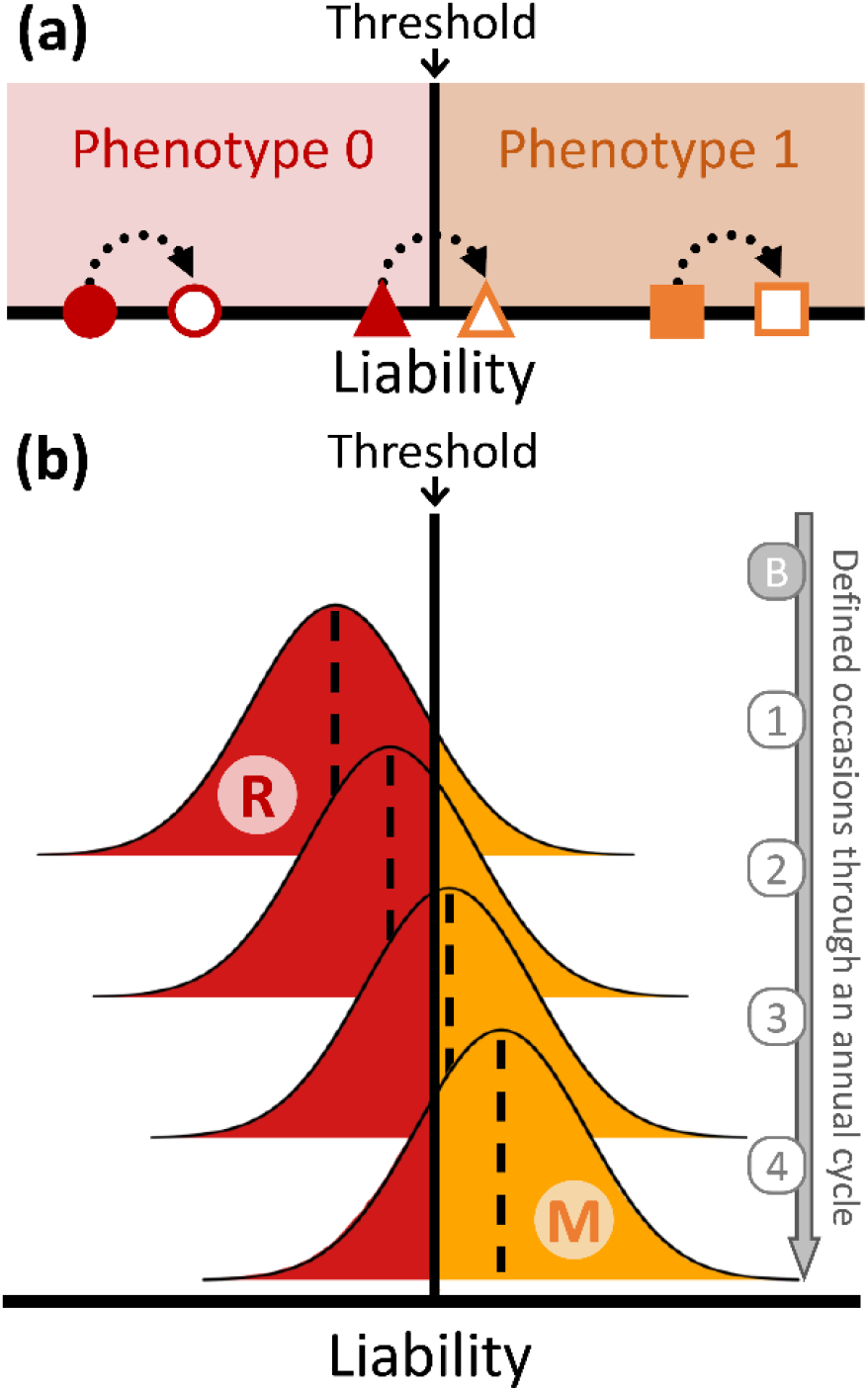
Multiscale principles of threshold trait variation. (a) Phenotype 0 versus 1 is expressed when the underlying continuous liability (x-axis) is below versus above a certain threshold. Filled symbols (circle, triangle, square) illustrate three individual liability intercepts of differing distance and direction from the threshold, reflecting genetic and/or permanent environmental effects. Open symbols represent current individual liabilities encompassing temporary environmental variation. Here, identical effects (dotted arrows) generate different degrees of within-individual phenotypic variation (the triangle individual is phenotypically plastic while the circle and square individuals are not), implying interactions between liability intercepts and temporary environments that generate among-individual variation in phenotypic plasticity. (b) The distribution of liabilities can change through time, illustrated here as a steady increase in mean liability (dashed vertical line) through an annual cycle of phenotypic expression for the trait seasonal migration (‘M’, orange) vs. residence (‘R’, red). In the breeding season (‘B’), there is no expression of M (i.e. all individuals express R) and hence no liability is represented. The liability is defined and represented on discrete occasions 1–4 through the non-breeding season. Changes in population mean on the liability scale generate changes in the phenotypic proportion of residents and migrants (red and orange areas below and above the threshold, respectively), and hence in both mean and variance on the phenotypic scale.

This threshold trait framework implies that phenotypic variation intrinsically results from emerging interactions among underlying additive genetic, permanent individual and temporary environmental effects. For example, the phenotypic effect of a particular allelic substitution will depend on the values of permanent and temporary environmental deviations. Hence, there can effectively be gene-by-environment interactions (‘G×E’) affecting the phenotype, even without any G×E acting directly on the liability scale (Figure 1; Reid and Acker 2022). Comprehensive dissection of interacting genetic and environmental effects on discrete traits therefore requires explicit consideration of all components of liability-scale variance and their transformation into phenotypic variances (as initiated by de Villemereuil et al. 2016; Acker et al. *in prep*). Such multi-scale variance decompositions are now required to fully understand and predict joint plastic and micro-evolutionary responses to environmental perturbations and changes.

The intrinsic conceptual complexities of discrete trait dynamics come hand-in-hand with substantial challenges of estimation. Observed dichotomous phenotypes must be used to estimate latent liability-scale variance components, which must then be back-transformed to derive phenotypic-scale variances. This is achievable using generalised linear mixed models (‘GLMMs’), where a non-linear data scale maps onto a linear latent scale through a link function (Nelder and Wedderburn 1972). By incorporating pedigree data describing genealogical relationships among individuals, such models can explicitly estimate latent-scale additive genetic variances, forming ‘animal models’ (Falconer and McKay 1996; Lynch and Walsh 1998; de Villemereuil et al. 2016). Dynamics of latent liabilities and resulting phenotypes within and across years and generations can then be estimated, allowing inference on how changing additive genetic and environmental effects on liabilities interactively translate into changing frequencies of dichotomous phenotypes.

Further major challenges arise in the common situation where phenotypic data are non-randomly missing from collected datasets, which can substantially bias quantitative genetic estimates and conclusions unless the (imperfect) observation process is modelled (Hadfield 2008; Nakagawa and Freckleton 2008; Steinsland et al. 2014). This is in principle achievable using joint capture-recapture and animal models (‘CRAMs’). But, to date, such models have only been developed for estimating additive genetic variance in survival, and implemented for one illustrative example (Papaïx et al. 2010; Morrissey et al. 2014; de Villemereuil 2018). General CRAMs, allowing estimation of variances underlying expression of any trait of interest given incomplete observation, would therefore substantially advance capabilities to dissect complex trait dynamics in the wild.

A prime example of these complexities of concept and estimation is seasonal migration versus residence, a critical life-history trait that directly shapes population dynamic outcomes in mobile animals inhabiting seasonally varying environments. Here, expression of seasonal migration (hereafter ‘migration’) can vary within sympatric-breeding populations, whereby some individuals remain resident at the breeding location year-round while other individuals migrate away for all or part of the non-breeding season before returning to breed. Resulting ‘partial migration’ occurs in numerous taxa, spanning fish, mammals, amphibians, reptiles, and birds (Lundberg 1988; Grayson et al. 2011; Chapman et al. 2012; Berg et al. 2019; Buchan et al. 2020; Chapman et al. 2011). Such phenotypic variation determines individuals’ seasonal locations and resulting exposure to heterogeneous seasonal environmental conditions, thereby directly affecting fitness, driving selection, and shaping spatio-seasonal population dynamics (Reid et al. 2018). Given a compound additive genetic and environmental basis, expression of migration could therefore directly mediate eco-evolutionary dynamics in seasonally varying environments.

Migration versus residence has long been broadly conceptualised as a quantitative genetic threshold trait (Berthold 1988; Pulido et al. 1996; Dodson et al. 2013). Substantial evidence shows that expression of migration, and closely related traits such as maturation or anadromy in salmonids, is highly polygenic (Pedersen et al. 2013; Lemopoulos et al. 2019; Sinclair-Waters et al. 2020; Bossu et al. 2022) and also environmentally dependent (affected by e.g. individual state, density, predation, weather, or food; Brodersen et al. 2008; Grayson et al. 2011; Eggeman et al. 2016; Yackulic et al. 2017; Boyle et al. 2010). Further, while migration is commonly highly phenotypically repeatable in iteroparous adults (e.g. Kerr et al. 2009; Grist et al. 2014; Zúñiga et al. 2017; Sawyer et al. 2018; Lehnert et al. 2018), considerable individual plasticity is also evident. Individuals can switch between migration and residence between winters (e.g. Grayson et al. 2011; Brodersen et al. 2014; Hegemann et al. 2015; Spitz et al. 2018; Xu et al. 2021) and within winters (representing late departure from or early return to breeding locations; Cagnacci et al. 2011; Fudickar et al. 2013; Reid et al. 2020). Such combinations of high repeatability and labile plasticity across nested timescales imply substantial variation in permanent environmental and/or genetic effects, and non-negligible temporary environmental effects. However, corresponding variances in liability for migration versus residence, and the degree to which such effects change through time and interactively translate into phenotypic change, have never been explicitly quantified in free-living natural populations. Consequently, we cannot yet predict the complex micro-evolutionary responses of partially migratory populations to environmentally-induced selection, or therefore predict the potential for evolutionary rescue through changing seasonal movement.

Progress requires phenotypic observations of migration versus residence in numerous individuals of known relatedness, within and across years and generations. While this can be achieved through large-scale field resightings of marked individuals, resulting datasets inevitably include heterogeneous detection failure due to spatiotemporal variation in observation effort and success (Acker et al. 2021a). Novel CRAMs that can partition variation in liability for migration while explicitly accounting for imperfect observation processes are therefore required.

Here, we provide a general CRAM that allows inference on liability-scale effects and variances for any dichotomous trait using individual capture-recapture histories. We fit this CRAM to 12 years of large-scale year-round individual resighting data that inform on individual migration versus residence, coupled with >30 years of accumulated pedigree data from a partially migratory population of European shags (*Gulosus aristotelis*). We draw three levels of inference. First, we estimate the components of liability-scale variance attributable to additive genetic and permanent individual versus temporary residual effects, and thereby evaluate the underlying potential for micro-evolution of seasonal migration. Second, we estimate the components of phenotypic-scale variance resulting from the non-linear genotype-environment-phenotype map that is intrinsic to the threshold trait model, thereby revealing how underlying liability-scale genetic and environmental variances interact to generate observed phenotypic variance. Third, we estimate changes in population-level mean additive genetic values, liabilities and resulting phenotypic proportions of migrants versus residents within and across the 12-year study, thereby revealing the expected magnitude and basis of temporal phenotypic dynamics of migration. Specifically, we examine the degree to which two known episodes of strong survival selection against residence induced by extreme climatic events (‘ECEs’; Acker et al. 2021a) caused persistent versus transient change in liability for migration, and in resulting phenotypic expression, through immediate micro-evolution versus changing distributions of permanent individual effects. Overall, we thereby provide new conceptual and empirical insights into how latent and expressed components of trait variation can shape micro-evolutionary and phenotypic responses in a critical life-history trait, seasonal migration versus residence, and thereby reshape spatio-seasonal population dynamics following environmental perturbations.

## METHODS

### General principles of the CRAM

Typically, on any single survey in a wild population study, some individuals are unobserved and consequently their phenotype cannot be measured (Lebreton et al. 1992; Nakagawa and Freckleton 2008). Sampling bias occurs when such observation failure is non-random with respect to phenotype (e.g. Biro and Dingemanse 2009). Yet, indirect inference on missing phenotypic data can still be made by modelling phenotype-dependent observation failure alongside the biological process of phenotypic variation. Further, while instances of observation failure can be evident in longitudinal individual-based datasets if individuals reappear after previous absences, observation failure cannot be directly distinguished from death or permanent emigration (i.e. apparent mortality) after an individual is last encountered. Consequently, apparent mortality must also be modelled as part of the biological process that shapes true phenotypic variation in the sampled population. These objectives can be achieved using capture-recapture models, i.e. state-space models representing individual transitions between alive and dead states across time parameterised by survival probability (‘*ϕ*’), and corresponding observations parameterised by detection probability (‘*p*’; Lebreton et al. 1992). Further, different ‘alive’ states can be defined to represent different phenotypes of interest, with within-individual phenotypic variation parameterised by state-transition probabilities (‘*ψ*’) that are conditional on survival. This generates a multistate capture-recapture model (Lebreton and Pradel 2002), which can be formulated to represent phenotypic variation in any discrete labile trait alongside phenotype-dependent survival and detection. Individual variation in *ψ* (and/or *ϕ*) can then be linked to a quantitative genetic animal model to form a CRAM, enabling estimation of components of (genetic) variance underlying phenotypic expression.

Non-random observation failure is inevitable when phenotypic data on migration versus residence are collected through field resightings of marked individuals in partially migratory populations (e.g. Grayson et al. 2011; Zúñiga et al. 2017), unless all potential non-breeding season locations are similarly observed. To handle this scenario, and hence estimate key components of underlying variance, we formulated a discrete-time multistate CRAM (Figure 2a). Here, the probability *ψ*_*i*_ that alive individual *i* transitions to the migrant phenotypic state (‘M’) corresponds to its probability of phenotype expression (i.e. its phenotypic expectation). Consequently, 1 − *ψ*_*i*_ is the probability of expressing the resident phenotype (‘R’; Fig 2a). Let *z*_*it*_ denote the phenotype of individual *i* at time *t* (assigning 0 for R and 1 for M), then:

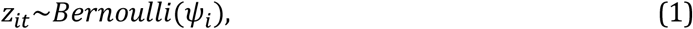

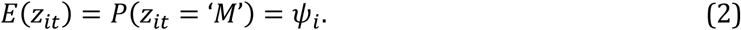

**Figure 2.**
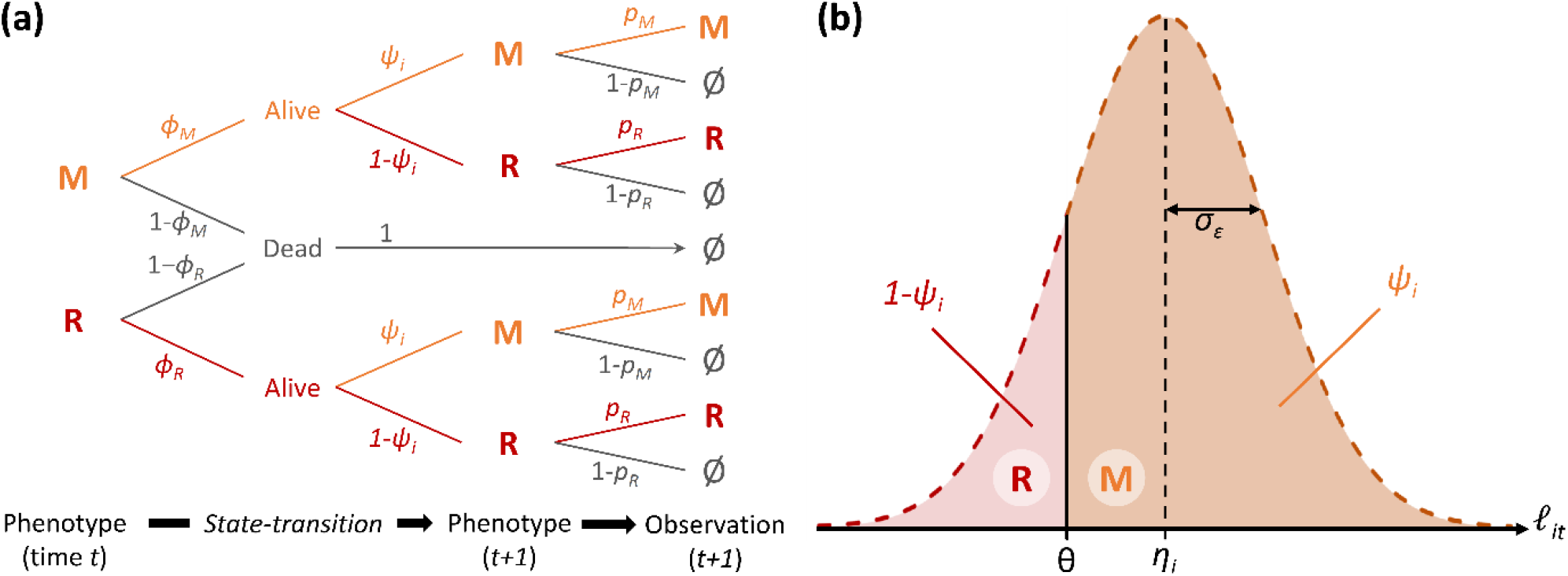
Discrete-time multistate capture-recapture model structure for a labile dichotomous threshold trait (e.g. seasonal migration versus residence). (a) Tree diagram summarising the probabilistic formulation of all possible outcomes for individual *i*, comprising transitions between phenotypic states from time *t* to *t* + 1, and observation at *t* + 1. Between time points, the individual either survives or dies. On any time point, a surviving individual is in phenotypic state ‘migrant’ (M) or ‘resident’ (R) and can be resighted in its current state (observation M or R respectively) or not resighted (Ø). Dead individuals cannot be resighted. Indices on the branches indicate the probability of corresponding steps along the focal path: *ϕ*_*M*_ and *ϕ*_*R*_ for survival of migrants and residents, *ψ*_*i*_ for expressing phenotype M; *p*_*M*_ for *p*_*R*_ for resighting of migrants and residents. For illustration purposes, survival and resighting probabilities are here represented simply as phenotype-dependent, but can be time- and location-dependent. (b) Illustration of how the individual phenotypic expectation *ψ*_*i*_ relates to the distribution of the time-specific individual liability *ℓ*_*it*_, assuming that *ℓ*_*it*_ is normally distributed with mean *η*_*i*_ and variance 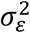. Under the threshold trait model, *ψ*_*i*_ is the probability that *ℓ*_*it*_ is greater than the threshold (θ). Accordingly, *ψ*_*i*_ is the area under the probability density function of *ℓ*_*it*_ between θ and +∞, which is the value at θ of the cumulative distribution function of a standard normal of mean 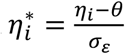. This is the probit transformation, *probit*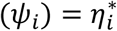.

Under the threshold trait model, *ψ*_*i*_ is the probability that the underlying continuous liability *ℓ*_*it*_ exceeds the threshold *θ*. Assuming that *ℓ*_*it*_ is normally distributed with mean *η*_*i*_ and variance 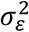 (Figure 2b):

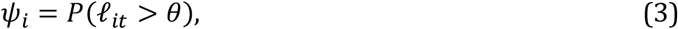

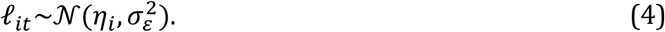

To decompose variance in liability to migrate into additive genetic and other permanent and temporary components we define an animal model on *ℓ*_*it*_, specified as a linear regression:

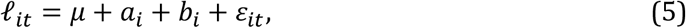

where *μ* + *a*_*i*_ + *b*_*i*_ = *η*_*i*_, *μ* is the overall mean, *a*_*i*_ is the additive genetic effect (‘breeding value’; 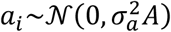, where A is the additive genetic relatedness matrix), *b* is the permanent individual effect (representing permanent environmental and non-additive genetic effects; 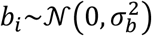, and *ε*_*it*_ is the temporary residual (temporary individual effect encompassing labile plasticity; 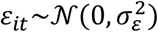. However, this animal model is non-identifiable because *ℓ* is a latent variable of unknown unit which cannot be measured. Since *θ* is therefore also unknown, the link to phenotypic inference on *ψ*_*i*_ from the capture-recapture model cannot be made directly through equation 3. Yet, these problems can be circumvented by considering a standardized model, where liabilities are threshold-centred and scaled by *σ*_*ε*_ (Dempster and Lerner 1950):

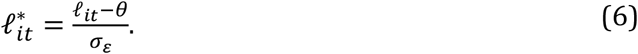

Standardised liability values 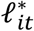 are therefore distances to the threshold and their unit is the standard deviation of the temporary residuals. This defines a reparameterised CRAM (equivalent to equations 1 and 2) where the underlying variable is 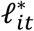, the threshold is 0, and the residual variance is 1:

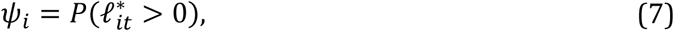

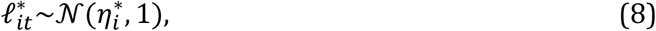

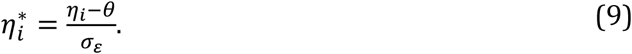

The phenotypic expectation *ψ*_*i*_ is therefore a simple function of 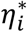, specifically 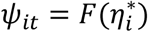, where F is the cumulative distribution function of the standard normal distribution. Since *F*^−1^ is the probit function, we retrieve a binomial GLMM with a probit link (de Villemereuil et al. 2016):

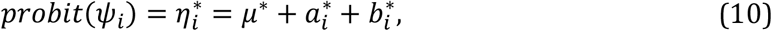

where 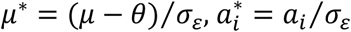, and 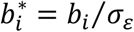.

Accordingly, we directly fit an identifiable animal model (equation 10) on the probit-transformed individual phenotypic expectation inferred by the capture-recapture model, and hence formulate the joint likelihood of the multistate CRAM. Here, it is critical to note that we estimate linear effects on the individual expectation 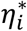 of the standardised liability for migration 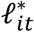, which is centred on *θ* and scaled by *σ*_*ε*_. Estimates of additive genetic and permanent individual variance in liability are therefore relative (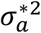 and 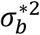), and scaled by the temporary residual variance (i.e. 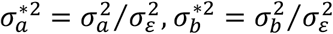).

### Study system and data collection

To quantify components of variance underlying migration versus residence using our CRAM, we collected intensive observations of non-breeding season locations of numerous pedigree-linked individuals in a partially-migratory shag population that breeds on Isle of May National Nature Reserve (hereafter ‘IoM’), Scotland (56°11′N, 2°33′W; Frederiksen et al. 2008; Daunt et al. 2014; Keogan et al. 2021).

During 1997–2021, ∼20600 chicks and ∼1100 adults were individually marked with a uniquely coded metal ring and an inscribed colour ring field-readable from ∼150m with a telescope. During non-breeding seasons (‘winters’) 2009–2021, we collected resightings throughout September–February at coastal sites where individuals roost daily to dry their partially-wettable plumage (Grist et al. 2014; Acker et al. 2021a; Supporting Information S1). We undertook ∼biweekly surveys of main roost locations on IoM and across the population’s migratory range in eastern Scotland, and also obtained occasional resightings from other locations spanning ∼800km of coast. When an individual was resighted, we assigned its phenotype as resident (if seen on or near IoM) or migrant (if seen elsewhere; Supporting Information S1).

During breeding seasons (April–June, ‘summers’), intensive monitoring of all nests and adjacent roost sites on IoM generated very high overall summer resighting probabilities of adults (0.90–0.98, mean 0.95 across 2010–2018), including identification of most ringed nest owners (∼95% of all breeders in 2009–2021; Acker et al. 2021a,b). Most adults were sexed through vocalizations and/or genotyping (Acker et al. 2021a). Hence, since shags are socially monogamous, putative mothers and/or fathers were assigned to ringed chicks.

Previous phenotypic analyses demonstrated structured and directional patterns of among- and within-individual variation in migration versus residence in shags (Reid et al. 2020; Acker et al. 2023), that are highly consistent with expectations for a threshold trait and inconsistent with simple Mendelian inheritance. This implies manifold genetic and/or environmental influences on migration, and hence that quantitative genetic variance partitioning is an appropriate route to inferring micro-evolutionary and phenotypic dynamics.

### Capture-recapture model design and resighting dataset

To infer individual phenotypic variation in migration versus residence, we built the multistate capture-recapture part of the CRAM using a full-annual-cycle structure previously devised for estimating survival selection (Acker et al. 2021a). Specifically, to maximise use of the year-round field resightings, we divided each winter into four ‘occasions’ on which individuals can express either phenotype M or R, and also included a summer occasion (Figure 1b, 2a). Although all focal individuals were assumed to be located on IoM in summer, meaning that summer sightings provide no information on winter locations or hence migration, utilising the summer resightings facilitates robust and precise inference on overall annual survival probabilities. This in turn facilitates inference on phenotypic variation and phenotype-dependent survival through intervening winters.

We defined six possible states that allowed us to represent the two phenotypes, M and R, while accounting for phenotype-dependent observation failure caused by spatio-temporal variation in resighting probability (Acker 2021a; Supporting Information S1). Specifically, the migrant phenotype was represented by five states corresponding to four geographically distinct migratory areas with non-zero but potentially differing resighting effort, plus a ‘ghost area’ representing all migrant destinations that were not surveyed (Acker et al. 2021a; Supporting Information S1). The resident phenotype was represented by one state corresponding to the IoM area.

Focal individuals enter the dataset during the first summer in which they were observed to breed on IoM during 2009–2020. In summer, all surviving individuals express phenotype R with probability 1. Between any two successive occasions across years, individuals either survive or die according to phenotype×sex×occasion×year-dependent (‘×’ denoting interactions) survival probability (*ϕ*). On each winter occasion, alive individuals express phenotype M or R according to probability *ψ*_*i*_ (where sources of variation are specified by the linear regression of probit(*ψ*_*i*_) constituting the animal model part of the CRAM described below). Conditional on expressing phenotype M, individuals go to one of the five possible migrant areas. If an individual’s phenotype was R in the previous occasion, it makes an initial move according to occasion×year×area-dependent destination probability *δ* (Supporting Information S2). If the individual’s phenotype was already M, it can go to another migrant area according to movement probability *γ*, which is low in our system, and hence was validly assumed constant across areas, occasions, and years (Acker et al. 2021a; Supporting Information S2). Alive individuals can then be resighted where they are, and hence directly observed as M or R, or not resighted, according to sex×occasion×year×area-dependent resighting (detection) probability *p* (fixed to zero in the ghost area and for dead individuals; Supporting Information S2).

To fit this model to our resighting data, we compiled individual capture-recapture histories of 2576 adult shags (1319 females, 1257 males) from 61281 year-round resightings (Supporting Information S1). Each history comprises a sequence of observation events coding whether and in which defined area an individual was resighted, spanning all occasions until summer 2021. Each history ended with an additional final observation event indicating whether the individual was resighted again between summer 2021 and mid-December 2021. Including this last datum avoided parameter redundancy and allowed independent estimation of *ϕ* and *p* across all focal time steps through the 2009–2021 study period (Supporting Information S2).

### Animal model design and pedigree dataset

We formulated the animal model part of the CRAM by considering the standardised individual liability expectation 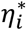, i.e. the probit-transformed probability *ψ*_*i*_ of expressing the migrant phenotype, thereby linking inference on trait variation with field observations analysed by the capture-recapture part of the CRAM (see above). We allowed overall mean liability to differ among the four defined annual winter occasions, representing seasonal progression in liability to migrate within each winter (e.g. Figure 1b). We also allowed mean liability to differ between females and males on each occasion. While current data are insufficient to estimate sex-specific additive genetic variance, modelling sex-specific means minimises the risk that variance estimates are inflated by sexual dimorphism. Further, to achieve our current ambition of providing first estimates of main components of liability- and phenotypic-scale variation in migration versus residence, we kept the animal model relatively simple by modelling constant additive genetic and permanent environmental variances in liability for all winter occasions. This effectively assumes an across-occasion genetic correlation of 1 (e.g. Figure 1b). We therefore specified:

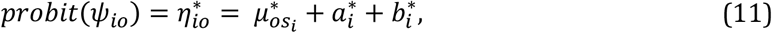

where parameters are as in equation 10, index *s*_*i*_ denotes the sex of individual *i*, and *o* denotes the winter occasion (nested within years; Supporting Information S2). Any further variation in liability occurring across the whole range of environmental variation experienced by individuals throughout the study period (i.e. within and among years) was thus considered as random within individuals (i.e. *ε*_*it*_ on the liability scale), representing the temporary residual variance which gives the unit of the standardised liability scale. We also fitted additional models that included effects of either brood, maternal, or paternal identity, to check whether shared developmental micro-environments caused resemblance among close kin that could have inflated estimates of 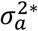. However, these additional variance components were negligible, and estimates of 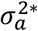 remained similar (Supporting Information S2).

Beyond model design, pedigree-based quantitative genetic inference fundamentally depends on pedigree structure. To calculate the A matrix containing the pairwise correlations between individual breeding values *a*_*i*_, we compiled pedigree data using observed social parentage of all phenotype-informative individuals (i.e. the 2576 individuals in the capture-resighting dataset) and their ancestors (Supporting Information S1). When same-brood individuals had an unringed parent, they were assigned a common dummy parent, thereby linking known siblings (e.g. Husby et al. 2010). However, 20% of phenotype-informative individuals (518) were not known to be related to any other phenotype-informative individuals. Accordingly, they provide little information for estimating 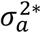, and any attempt to estimate their breeding values 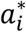 would be biased towards their phenotype which is also influenced by environmental effects (Postma 2006; Hadfield 2010). Hence, for these individuals, we replaced the animal model part of the CRAM with a simpler regression that did not distinguish additive genetic from permanent individual effects and instead simply considered their sum (Supporting Information S2):

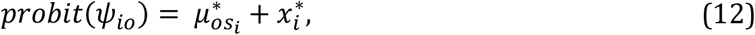

where 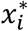 is the total individual effect 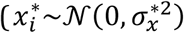 and 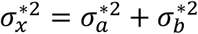. Accordingly, the pedigree utilised to derive relatedness values that informed estimation of 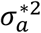 comprised 2349 individuals, of which 88% (2058) were phenotype-informative and 12% (291) were additional ancestors (including 86 dummy parents). Overall, 29% of these phenotype-informative individuals (602) and 90% of their additional ancestors (262) had both parents unknown and no siblings in the pedigree, and hence comprised the defined founder population of (presumed) unrelated individuals for which 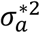 is estimated (Kruuk 2004; Wolak and Reid 2017). All other individuals (63% of the utilized pedigree, 1485 individuals) were descendants of the founders and had both parents identified with known or dummy identities, hence achieving relatively high pedigree completeness (Wolak and Reid 2017).

From this pedigree, we computed all pairwise additive genetic relatedness values (i.e. twice the expected coefficient of co-ancestry or kinship) between phenotype-informative individuals, i.e. the A matrix elements (Supporting Information S1). There were 8618 pairwise non-zero values, of which 18% were ≤0.0625, 20% were 0.125, 31% were 0.25, and 31% were 0.5 (mean and median 0.25, standard deviation 0.17). Phenotype-informative individuals had a mean of 8.4 links with others (range 1–48, median 6, standard deviation 7.4). The per-individual mean of non-zero relatedness values was 0.35 (range 0.06–0.50, standard deviation 0.12) with a prominent mode at 0.5. Overall, this distribution demonstrates the existence of numerous close pedigree links, providing valuable information for estimating 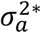.

### Analyses of the multistate CRAM

We coded our multistate CRAM in Stan, a probabilistic programming language for Bayesian inference, using package rstan in R to sample the posterior distribution of each parameter (Carpenter et al. 2017; R core team 2022). For the capture-recapture model parameters, comprising probabilities, we used uniform priors (Supporting Information S2). We retrieved estimates of survival, movement, and detection probabilities that were consistent with those obtained from our previous foundational models that did not include an animal model or estimate liability-scale parameters (Acker et al. 2021a, b). For the animal model parameters, we used weakly informative priors, and sensitivity analyses showed that conclusions were robust to different prior choices (Supporting Information S2). A ‘control’ CRAM, fitted after randomizing individual identities in the A matrix, showed no bias in estimated 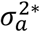 (Supporting Information S2). Posterior predictive checks (Gelman et al. 1996) devised for the multistate capture-recapture model (Acker et al. 2021a) indicated good overall fit. Details of posterior sampling procedures, diagnostics, and all parameter estimates are in Supporting Information S4. Complete numerical summaries and posterior samples of all parameters are archived in Zenodo alongside full Stan and R codes (see *Data accessibility statement*). Hereafter, estimates are presented as posterior means with 95% credible intervals (‘95%CI’).

### Derived calculations

To achieve full liability-scale and phenotypic-scale variance decompositions for migration versus residence, we derived posterior distributions of compound and transformed quantities from the posterior samples of the CRAM parameters. First, we extracted the total variance 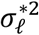 in liability on the standardised scale (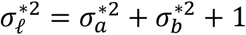, where 1 is the temporary residual variance 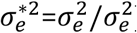). We then calculated the proportion of the total variance in liability (conditional on the fixed annual progression through winter occasions) attributable to the total individual variance (i.e. repeatability, 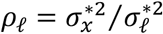) and to additive genetic variance (i.e. narrow-sense heritability, 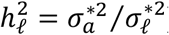).

Second, to quantify phenotypic-scale variation resulting from the estimated liability-scale variation, we derived the overall phenotypic mean 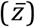 and phenotypic variance (*V*_*z*_; where *V* denotes phenotypic-scale variances, while *σ*^2^ denotes liability-scale variances). We then partitioned *V*_*z*_ into phenotypic-scale additive genetic variance (*V*_*A*_) versus other components that involve permanent individual or temporary environmental variances (mathematical expressions for back-transformation in Supporting Information S3; Acker et al. *in prep*). Given general properties of non-linear mappings of dichotomous variables to underlying continuous variables, the phenotypic mean and variance become interdependent (Figure 1b). Accordingly, our model implies occasion×sex-specific 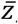, and hence *V*_*z*_ (and its components). In addition, because some variation that is strictly additive on the liability-scale becomes non-additive on the phenotypic-scale (Figure 1a, Dempster and Lerner 1950; de Villemereuil et al. 2016), new components of phenotypic variance emerge from interactions between effects that are modelled as additive on the liability scale. More precisely, *V*_*z*_ is the sum of phenotypic variances resulting independently from liability-scale additive genetic effects, permanent individual effects, and temporary residual effects (respectively *V*_*a*_, *V*_*b*_ and *V*_*ε*_) plus phenotypic variances resulting from all possible interactions (i.e. *V*_*a*×*b*_, *V*_*a*×*ε*_, *V*_*b*×*ε*_, and *V*_*a*×*b*×*ε*_; Supporting Information S3). Similarly, interactions between genes underpinning the liability-scale additive genetic effects emerge on the phenotypic scale, implying that *V*_*A*_ is a subcomponent of *V*_*a*_, alongside emerging non-additive genetic variance *V*_*NA*_ (Dempster and Lerner 1950; Robertson 1950; de Villemereuil et al. 2016; Supporting Information S3). From this decomposition, we further derived the phenotypic-scale total individual variance (*V*_*x*_ = *V*_*a*_ + *V*_*b*_ + *V*_*a*×*b*_), repeatability (*ρ*_*z*_ = *V*_*x*_/*V*_*z*_), and narrow-sense heritability 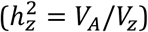.

Finally, we used our CRAM inferences to derive estimates of population-level dynamics of breeding values, permanent individual values and overall liabilities for migration and resulting phenotypic expression through the 12 study years, thereby revealing micro-evolutionary and phenotypic changes. Specifically, we derived the mean breeding value (*ā*^*^), mean permanent individual effect 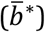, mean standardised liability expectation 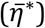, and expected proportion of migrants 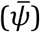 for each occasion in each year (Supporting Information S3). Here, temporal changes in *ā*^*^ and 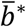 (and hence in 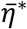 and 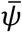) result from mortality of individuals already present in the focal adult population, and from entry of new individuals every summer. When individuals were never seen again after an episode of selection, their contributions to changes in *ā*^*^ and 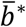 were inferred as functions of survival probabilities, which are phenotype-dependent in our CRAM.

Estimated changes in *ā*^*^ and 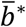 thus partly rely on the assumption that individuals’ current phenotypes are the sole cause of survival selection (Morrissey et al. 2010, 2012; Supporting Information S3). We did not explicitly quantify overall trends in *ā*^*^ or 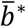 across the 12 years, but focused on evaluating the evidence for changes (‘Δ’) in *ā*^*^ or 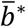 between two consecutive time points surrounding the two known episodes of strong survival selection against residence (in late-winter 2012-13 and 2017-18; Acker et al. 2021a). We computed posterior distributions of Δ and calculated the probability ‘*P*_*Δ*_’ that Δ had the same sign as its posterior mean (Supporting Information S3). *P*_*Δ*_ ≈1 indicates strong evidence for a positive or negative change, while *P*_*Δ*_ ≈0.5 indicates no clear evidence for either.

## RESULTS

### Seasonal change in mean liability and phenotypic expression of migration

Our model retrieved expected patterns of seasonal change in mean liability to migrate, and hence in phenotypic expression of migration versus residence (i.e. in overall phenotypic mean 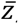). Specifically, the overall liability-scale intercept *μ* was lowest and negative in September (winter occasion 1), then increased to values close to the threshold in October–February (winter occasions 2–4; Table 1, Figure 3). This general pattern of winter progression in liability was broadly similar in both sexes; *μ* was slightly lower in males than females in occasion 1, with no clear sex differences in occasions 2–4 (Table 1, Figure 3). Accordingly, the defined founder population was expected to contain more residents than migrants in September (occasion 1), particularly in males, and then similar proportions through the rest of the winter (i.e. occasions 2–4; Figure 3). The estimated liability-scale variation therefore implies substantial phenotypic-scale variation.

**Table 1.**
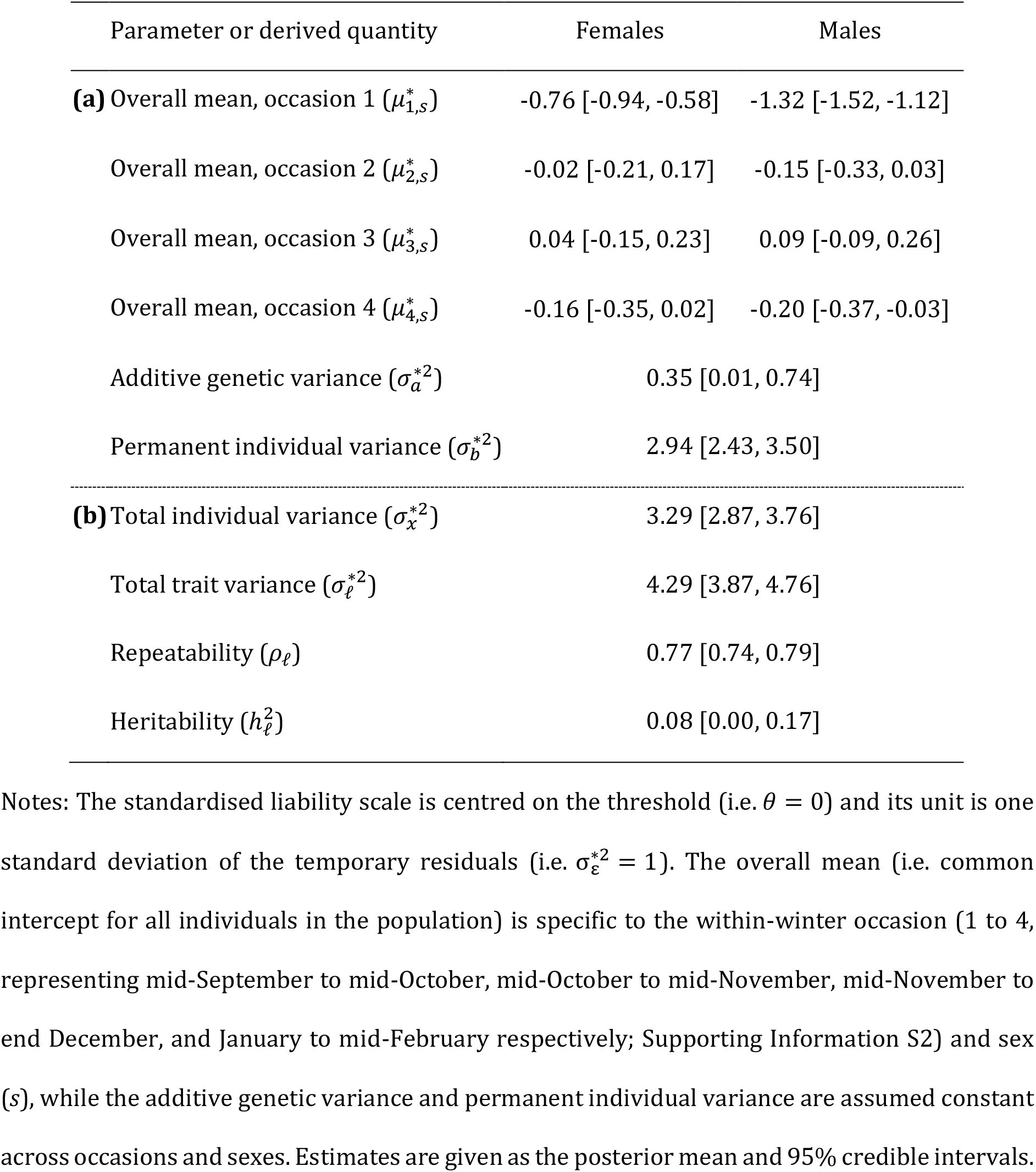
Capture-recapture animal model (CRAM) estimates of (a) sex-specific occasion effects and variance components of the standardised liability underlying seasonal migration versus residence, and (b) derived quantities.

| Parameter or derived quantity | Females | Males |
| --- | --- | --- |
| **(a)** Overall mean, occasion 1 ($\mu_{1,s}^*$) | -0.76 [-0.94, -0.58] | -1.32 [-1.52, -1.12] |
| Overall mean, occasion 2 ($\mu_{2,s}^*$) | -0.02 [-0.21, 0.17] | -0.15 [-0.33, 0.03] |
| Overall mean, occasion 3 ($\mu_{3,s}^*$) | 0.04 [-0.15, 0.23] | 0.09 [-0.09, 0.26] |
| Overall mean, occasion 4 ($\mu_{4,s}^*$) | -0.16 [-0.35, 0.02] | -0.20 [-0.37, -0.03] |
| Additive genetic variance ($\sigma_a^{*2}$) | 0.35 [0.01, 0.74] | |
| Permanent individual variance ($\sigma_b^{*2}$) | 2.94 [2.43, 3.50] | |
| **(b)** Total individual variance ($\sigma_x^{*2}$) | 3.29 [2.87, 3.76] | |
| Total trait variance ($\sigma_\ell^{*2}$) | 4.29 [3.87, 4.76] | |
| Repeatability ($\rho_\ell$) | 0.77 [0.74, 0.79] | |
| Heritability ($h_\ell^2$) | 0.08 [0.00, 0.17] | |

Notes: The standardised liability scale is centred on the threshold (i.e. $\theta = 0$ ) and its unit is one standard deviation of the temporary residuals (i.e. $\sigma_\varepsilon^{*2} = 1$ ). The overall mean (i.e. common intercept for all individuals in the population) is specific to the within-winter occasion (1 to 4, representing mid-September to mid-October, mid-October to mid-November, mid-November to end December, and January to mid-February respectively; Supporting Information S2) and sex ( $s$ ), while the additive genetic variance and permanent individual variance are assumed constant across occasions and sexes. Estimates are given as the posterior mean and 95% credible intervals.

**Figure 3.**
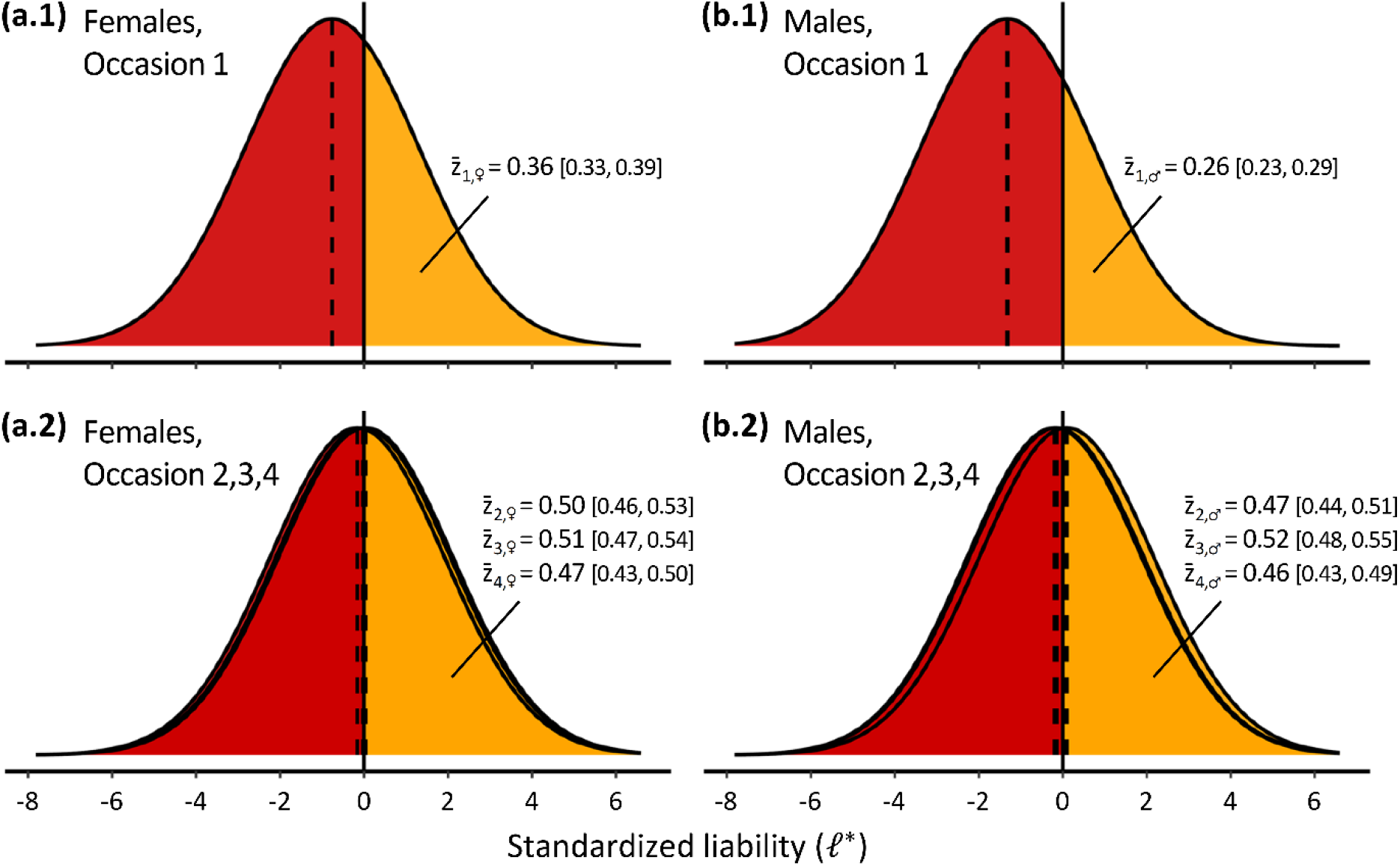
Density curves for occasion-specific distributions of the standardised liability to migrate (*ℓ*^*^) in (a,c) females and (b,d) males, for (a,b) seasonal occasion 1 (mid-September to mid-October) and (c,d) occasions 2-4 (spanning the rest of the winter), estimated from the capture-recapture animal model (CRAM). Dashed vertical lines represent distribution means (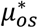 for any winter occasion o and sex s). Solid vertical lines represent the threshold (0 on the standardised liability scale). The area underneath the curve for *ℓ*^*^ > 0 (in orange) is the phenotypic mean 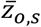, i.e. the expected proportion of migrants (and the area for *ℓ*^*^ < 0, in red, is the expected proportion of residents1 − 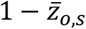). On c and d, curves for occasions 2-4 are superimposed because they are very similar. Numerical estimates of 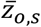 are shown as posterior mean and 95% credible intervals. Numerical estimates for 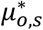 and 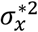 are in Table 1.

### Liability-scale variance decomposition

The posterior distributions of the additive genetic variance 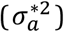 and permanent individual variance 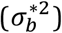 in liability for migration (on the standardised liability scale) showed peaks of density that were clearly away from zero and hence distinct from the prior distributions (Figure 4). The magnitude of 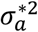 was non-negligible, corresponding to approximately one third of the temporary residual variance (Figure 4a, Table 1), and demonstrating potential for micro-evolutionary change in liability for migration in response to selection. Meanwhile, 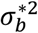 was notably large, corresponding to approximately three times the temporary residual variance (Figure 4b, Table 1). Consequently, there was very high liability-scale repeatability (*ρ*_*ℓ*_ ≈ 77%, Table 1), and modest liability-scale heritability (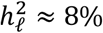, Table 1).

**Figure 4.**
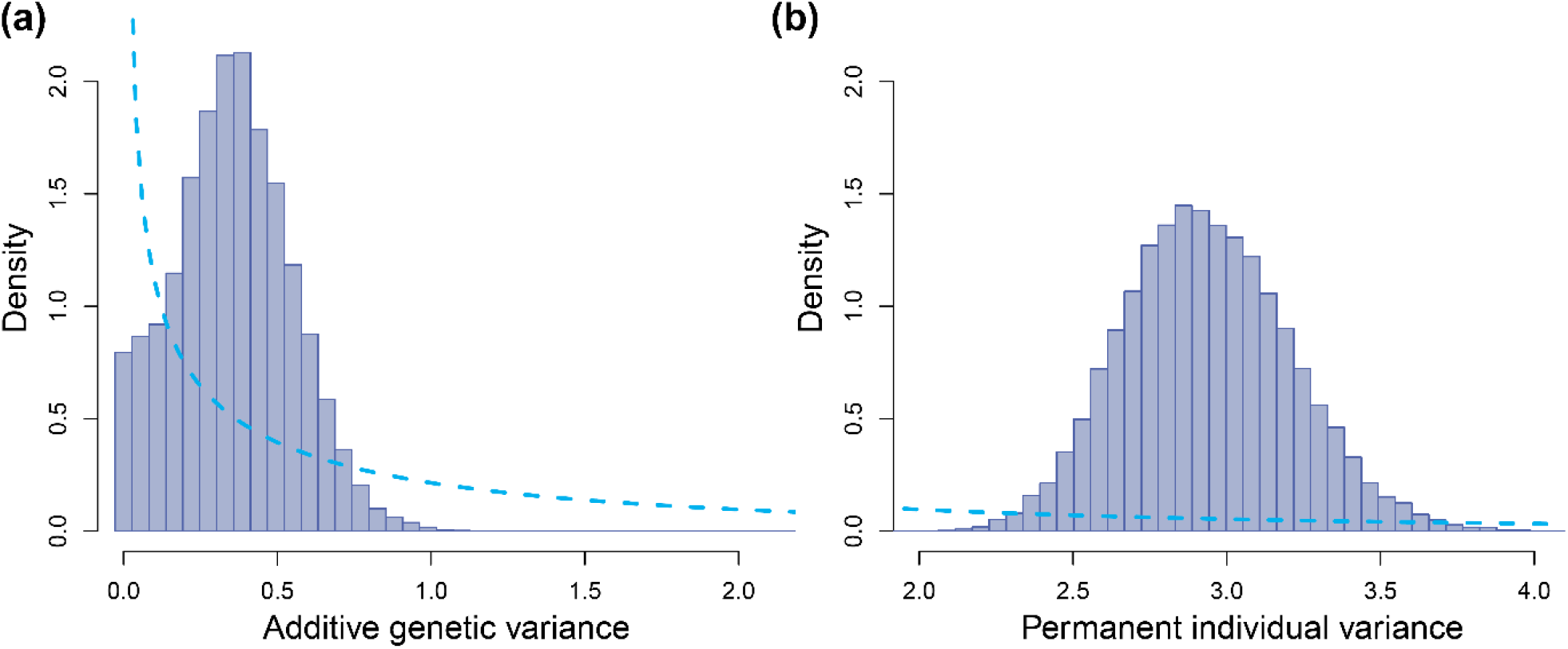
Prior (dotted line) and posterior (bars, MCMC samples) distributions of (a) additive genetic variance 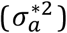 and (b) permanent individual variance 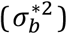 in liability for seasonal migration versus residence. Estimates are on the standardised liability scale, where the unit is one standard deviation of the temporary residuals of the liability (*σ*_*ε*_).

### Phenotypic-scale variance decomposition

Due to the large total variance in liability, and the proximity of the overall mean liability to the threshold, the total phenotypic variance in expression of migration versus residence *V*_*z*_ was always close to, or at, the maximum possible for a dichotomous trait (∼0.2–0.25; Figure 5). Transformations of variance components from the liability scale to the phenotypic scale showed that the largest component of *V*_*z*_ was the variance coming from liability-scale permanent individual effects (*b*) independently of other liability-scale effects (comprising ∼48% of *V*_*z*_; Figure 5). Meanwhile, the component of *V*_*z*_ coming purely from liability-scale additive genetic effects (*a*) was small, and essentially comprised phenotypic-scale additive genetic variance, yielding a phenotypic-scale heritability 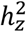 of ∼5% (Figure 5). Further, variance resulting from the emerging interaction between *a* and *b* constituted ∼2% of *V*_*z*_ (Figure 5). Consequently, the high overall phenotypic repeatability of seasonal migration versus residence (*ρ*_*z*_ ≈ 55%; Figure 5) is primarily due to permanent individual rather than additive genetic effects.

**Figure 5.**
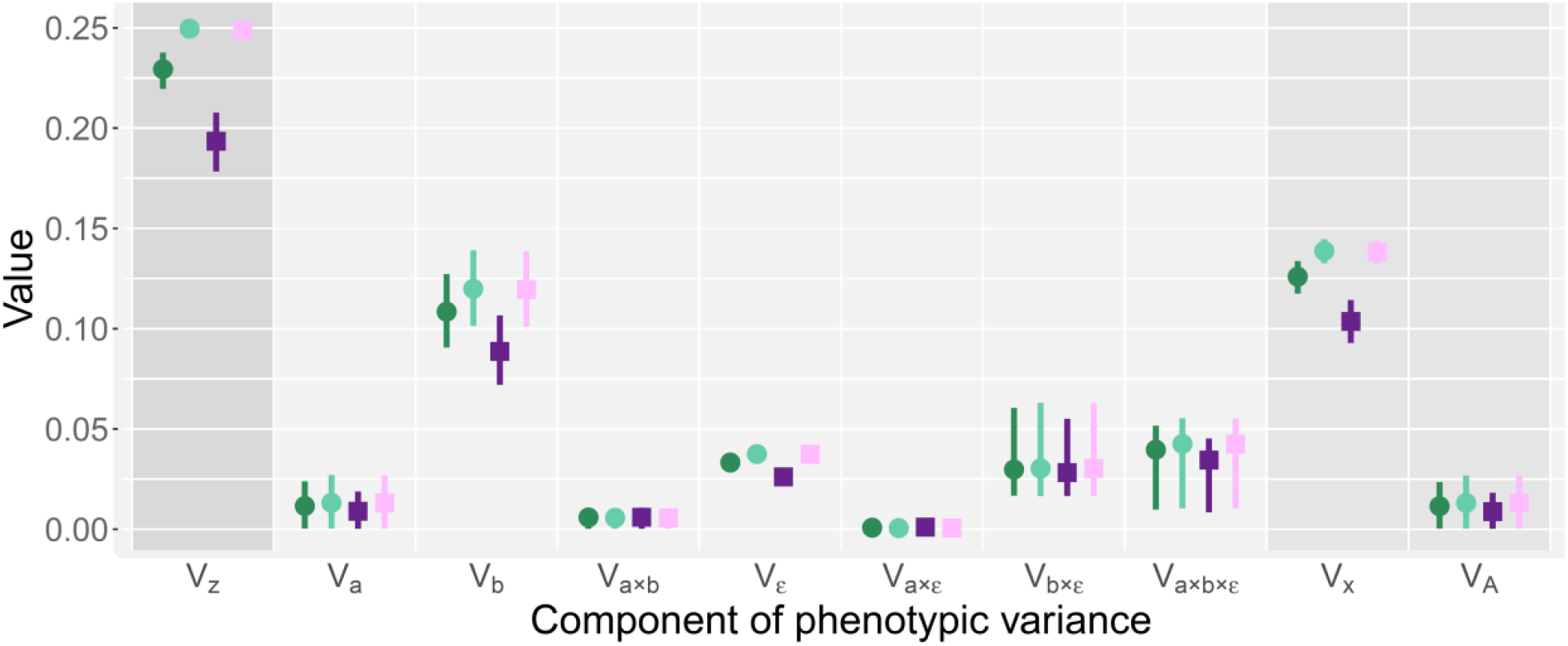
Decomposition of phenotypic variance (*V*_*z*_) in seasonal migration versus residence, for females (green circles) and males (purple squares) in winter occasion 1 (dark colouration, mid-September to mid-October) and occasion 2 (light colouration, mid-October to mid-November). Estimates for occasions 3 and 4 (spanning the rest of the winter) are virtually identical to occasion 2 (to two decimal places; Supporting Information S3). *V*_*z*_ is the total phenotypic variance. *V*_*a*_, *V*_*b*_ and *V*_*ε*_ are the phenotypic variances arising independently from liability-scale additive genetic effects *a*, permanent individual effects *b*, and temporary residual effects *ε*, respectively. *V*_*a*×*b*_, *V*_*a*×*ε*_, *V*_*b*×*ε*_, and *V*_*a*×*b*×*ε*_ are the phenotypic variances arising from interactions between these liability-scale effects. *V*_*x*_ is the total phenotypic-scale individual variance. *V*_*A*_ is the phenotypic-scale additive genetic variance. Point estimates are posterior means and lines indicate 95% credible intervals. Numerical details, including proportions of *V*_*z*_, are in Supporting Information S4.

Of the remaining 45% of *V*_*z*_, constituting temporary phenotypic variance, approximately one third came purely from liability-scale temporary residual effects *ε* (i.e. comprising ∼15% of *V*_*z*_). The rest, comprising ∼30% of *V*_*z*_, came from non-additive effects emerging from interactions between *ε, a* and *b*, encompassing among-individual variation in labile phenotypic plasticity (Figure 5). Here, the variance coming from *a* × *ε* was very small (<1% of *V*_*z*_), but those coming from *b* × *ε* and *a* × *b* × *ε* were substantial (∼12% and ∼17% respectively).

These outcomes were broadly similar for both sexes across all four defined winter occasions (Figure 5). The only notable quantitative difference was that the total phenotypic variance *V*_*z*_ was slightly lower in males than females in occasion 1 (i.e. mid-September to mid-October, Figure 5).

### Population-level changes across the study period

Model predictions of population-level changes in liability and phenotypic expression of migration across the study period revealed multiscale responses to known episodes of strong ECE-induced survival selection. Mean breeding value in the adult population (*ā*^*^) was stable across the first four years with posterior means slightly above the founder population mean of zero, then increased abruptly (to ∼0.03 on the standardised liability scale) following the first known episode of strong selection against residence during late-winter 2012-13 (Figure 6a). Subsequently, *ā*^*^progressively decreased (down to ∼0.01) until the next selection episode in late-winter 2017-18, when it increased abruptly (to ∼0.03), then remained stable across subsequent years (Figure 6a). There was high support for instantaneous positive changes in *ā*^*^ predictions following each selective ECE (*P*_*Δ*_ =0.92), indicating micro-evolutionary responses to selection manifested as increased mean breeding value in the adult population.

**Figure 6.**
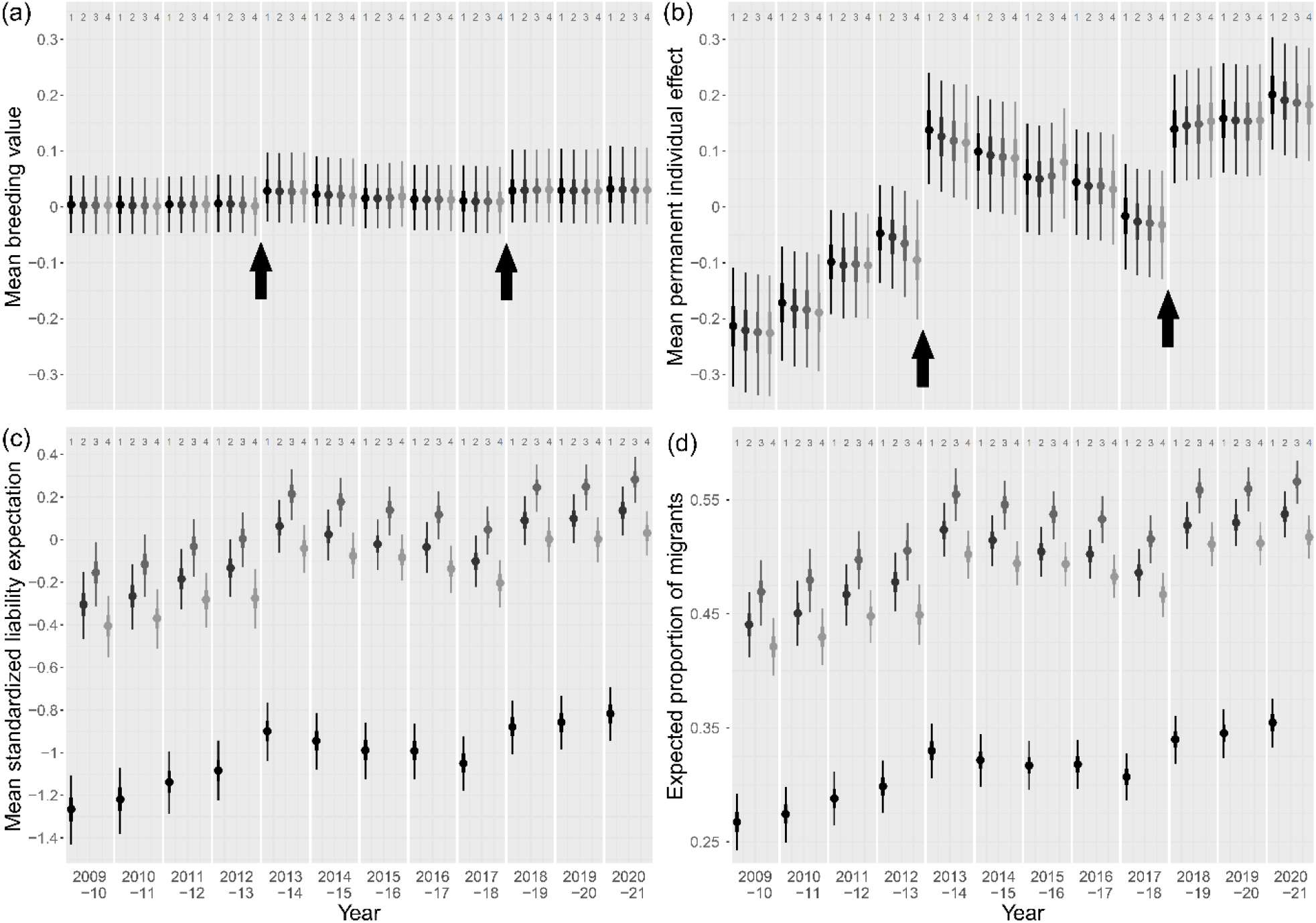
Estimated values of population means of (a) breeding values (*ā*), (b) permanent individual effects 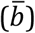, (c) standardised liability expectation 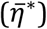 and (d) expected proportion of migrants 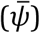, across the four defined occasions within each of the twelve focal biological years. The within-winter occasion (1 to 4) is indicated on top row and distinguishable across years by colouration shade (from black to light grey). Point estimates are posterior means, inner and outer line segments indicate 50% and 95% credible intervals. Black arrows on panels (a) and (b) point at the two episodes of strong survival selection that occurred in late-winter 2012-13 and 2017-18.

The mean permanent individual effect 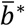 also increased dramatically following the two episodes of ECE-induced selection (Figure 6b, *P*_*Δ*_ =1), representing major shifts in the adult distribution of liabilities that are not heritable across generations. Further, 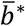 also varied between other consecutive years (Figure 6b; Supporting Information S4), largely reflecting variation in permanent individual effects among cohorts of recruiting adults.

Overall, the concurrent increases in *ā*^*^ and 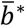 following the two selection episodes generated substantial increases in the mean standardised liability expectation 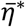 (Figure 6c), which in turn translated into moderate increases in the expected phenotypic proportion of migrants versus residents 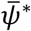 (Figure 6d). Due to the relative proximity to the threshold, changes in 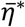 translated almost linearly to changes in 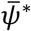 and primarily reflected within-year winter progression and between-year variation in 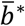 rather than variation in *ā*^*^ (Figure 6).

## DISCUSSION

Understanding population responses to environmentally-induced selection requires dissecting genetic and environmental components of variation in key traits; but these components become complex and entangled for phenotypically discrete traits. Our explicit decompositions of liability-scale and phenotypic-scale variation in the ecologically critical trait of seasonal migration versus residence demonstrate non-negligible additive genetic variance in liability and show how this variation is manifested phenotypically, both independently and in interaction with permanent and temporary environmental effects. Our results reveal the potential for both rapid micro-evolutionary change and phenotypic inertia in migration versus residence, as manifested in response to observed episodes of strong survival selection, and highlight the intrinsic emergence of complex interactions between micro-evolution and environmental variation in dichotomous threshold traits.

### Multiscale variance partitioning and micro-evolutionary potential

Our application of a novel multistate CRAM to full-annual-cycle field observations revealed non-trivial additive genetic variation in liability for seasonal migration in a wild partially migratory population, implying potential for micro-evolutionary responses to selection. In principle, the non-linear genotype-phenotype map that is fundamental to the threshold trait concept implies that liability-scale additive genetic variation could become partly non-additive on the phenotypic scale on which selection directly acts (Dempster and Lerner 1950; de Villemereuil et al. 2016). Intrinsic genetic interactions can therefore emerge and hide heritable liability-scale variation, generating ‘cryptic’ genetic variation which is apparently shielded from direct phenotypic selection (Roff 1996, 1998; Moorad and Linksvayer 2008; McGuian and Sgrò 2009; Pulido 2011). Yet, for threshold traits in constant environments, yielding constant fitness differences between the alternative phenotypes, the proportion of genetic variation that is cryptic is greater, and selection on genetic variation weaker, when the frequency of the commonest phenotype is higher (Falconer and McKay 1996; Roff 1998; Moorad and Linksvayer 2008). In our system, since frequencies of migration versus residence are close to parity (especially in winter occasions 2–4; Figure 5), most additive genetic variance estimated on the liability scale is phenotypically expressed. Accordingly, any selection acting on the migrant versus resident phenotypes should generate maximum selection intensity on liability, and hence maximum potential magnitude of micro-evolutionary response.

However, our decompositions and scale-transformations also highlight that intrinsic gene-by-environment interactions can arise in labile threshold traits, which has not previously been fully considered (Reid and Acker 2022; Acker et al. *in prep*). Specifically, we show that much of the total phenotypic variance in migration versus residence results from intrinsic interactions among liability-scale components, notably from a three-way interaction involving permanent individual and temporary environmental effects alongside additive genetic effects (Figure 5). This interaction emerges even though our current CRAM did not explicitly consider any gene-by-environment interactions in liability, as could be conceptualised through reaction norms of liability-scale plasticity. Instead, the phenotypic-scale interactions result from variation in total individual effects on liability, which generate among-individual variation in the degree and direction of labile phenotypic plasticity caused by temporary environmental deviations (Figure 1; Reid and Acker 2022; Acker et al. 2023). Such phenotypic plasticity may then be subject to selection, for example resulting from induced costs of phenotypic change, or benefits of rapid phenotype-environment matching (DeWitt et al. 1998; Auld et al. 2010; Murren et al. 2015). Indeed, we previously demonstrated episodes of selection on phenotypic plasticity in migration versus residence (manifested as within-winter switching) in shags, of magnitudes that varied among years (Reid et al. 2020; Acker et al. 2023). Major discrepancies in the form and/or magnitude of selection on liabilities, and hence in micro-evolutionary responses, could therefore arise compared to expectations from existing threshold-trait theory that only considers the relative frequencies and fitness of current alternative phenotypes (Falconer & Mackay 1996; Roff 1996, 1998; de Villemereuil et al. 2016). Meanwhile, the joint multiscale variation in liability and phenotypic plasticity in threshold traits also implies fundamentally different evolutionary dynamics of plasticity than expected for continuous traits where reaction norms directly act on observed phenotypic scales (Nussey et al. 2007; Lande 2009; Reid and Acker 2022).

Moreover, permanent individual variance was notably the largest component of liability-scale variance underlying migration versus residence in shags (Figure 4; Table 1), causing substantial phenotypic variance in isolation and through interactions (Figure 6). This variance component encompasses non-additive genetic effects acting on the liability scale and/or lasting effects of early-life environments that induce repeatable among-individual variation (Scheiner 1993; Kruuk 2004; Beldade et al. 2011). Variation in early-life conditions could therefore be one main determinant of population distributions of liabilities and resulting proportions of migrants versus residents, hence defining the extent of adult labile phenotypic plasticity and setting the stage for survival selection to further shape distributions of liabilities (Figure 1; Reid and Acker 2022). Plastic development could thereby fundamentally shape the combined potential for subsequent labile plastic and micro-evolutionary responses to environmental perturbations and changes, thereby shaping individual and population capacities to buffer environmental impacts (Beaman et al. 2016).

### Micro-evolutionary potential in partial migration and threshold traits

Despite the key role that rapid phenotypic change in migration versus residence could play in driving spatio-seasonal eco-evolutionary dynamics in diverse animals (Pulido 2011; Reid et al. 2018; Hsiung et al. 2018), few studies have explicitly quantified key components of genetic and environmental variation. Moreover, standardised measurement of evolutionary potential is particularly challenging for threshold traits, impeding cross-study comparisons. ‘Evolvability’, specifically defined as the mean-scaled additive genetic variance (Houle 1992; Hansen et al. 2011; Hansen and Pélabon 2021), is meaningless on the liability scale, because values are arbitrarily measured as deviations from the threshold, and additive genetic variance is only estimable relative to environmental variance. We are thus left with heritability as a standardized measure of additive genetic variation, which may primarily reflect the magnitude of environmental variance and hence have little value as a comparator of evolutionary potential (Houle 1992; Hansen et al. 2011; Hansen and Pélabon 2021). Arbitrarily high heritabilities can be expected in studies conducted under controlled laboratory conditions or across short time periods, which experience greatly restricted environmental variation.

Indeed, the very few previous studies that quantified additive genetic variance in liability for migration versus residence reported ∼50% heritability, contrasting with our 9% (Table 1). One study of brook trout (*Salvelinus fontinalis*) included only three years with two cohorts overlapping in one year, in brook trout (Thériault et al. 2007), and two others used captive bred and raised Atlantic salmon (*Salmo salar*; Páez et al. 2011; Debes et al. 2020). Similarly high heritabilities of proxies of seasonal migration were unsurprisingly reported by other studies conducted under controlled captive conditions (e.g. body size, body mass, or age at maturation in salmonids: Dodson et al. 2013; migratory activity in blackcaps, *Sylvia atricapilla*: Berthold and Pulido 1994; Pulido et al. 1996) and for threshold traits in general (Roff 1996). Moreover, most such previous studies focused on threshold traits where phenotypic expression is (approximately) fixed, and hence not subject to any temporary environmental variance. In fact, relatively small heritabilities have been reported in other labile threshold traits that can vary within individuals (e.g. survival, Krebs and Loeschke 1997, Papaïx et al. 2010; divorce, Germain et al. 2018). Our analytical and conceptual advances, which allow estimation and interpretation of variances underlying discrete traits from incomplete observations, should now facilitate new evaluations of components of liability-scale and phenotypic-scale variation in migration, and other threshold traits, in wild populations.

### Temporal dynamics and response to selection

Our dissection of the temporal dynamics of liabilities underlying migration versus residence reveal how phenotypic dynamics can emerge within and across full annual cycles. Notably, our analyses show detectable increases in mean breeding values following the two known episodes of strong phenotypic survival selection against residence associated with extreme late-winter storms in 2012-13 and 2017-18 (Figure 6a). Further, these estimates of micro-evolutionary responses are likely conservative, since our calculations partly assumed that dichotomous migrant versus resident phenotypes are the sole cause of selection on liability-scale breeding values (Supporting Information S3). In fact, there could be stronger directional selection along the phenological continuum of phenotypes ranging from full year-round residence to full-winter migration, which partly reflects additive genetic variation in liability (Acker et al. 2023). More generally, selection on liability could also take subtler forms than the simple direct selection on dichotomous phenotype that our CRAM currently assumes (including disruptive or stabilising selection; Acker et al. 2023; Reid and Acker 2022). The indications that ECE-induced selection caused immediate micro-evolutionary shifts in mean liability-scale breeding values for a key life-history trait that are large enough to be evident between consecutive seasonal timesteps in a wild population are therefore remarkable. In contrast, other individual-based wild population studies have commonly struggled to uncover (cross-generational) micro-evolutionary changes in traits that apparently experience strong directional selection spanning several years or decades (e.g. Gienapp et al. 2006; Bonnet et al. 2019; Biquet et al. 2022).

Nonetheless, the changes in population mean breeding values following the ECEs were dwarfed by simultaneous changes in mean permanent individual effects (Fig 6a,b, which may also be somewhat conservative estimates). Increases in phenotypic expression of migration versus residence following the ECEs therefore primarily reflect selection on permanent individual effects within the focal adult population rather than on additive genetic breeding values. Yet, the phenotypic inertia in increased migration that might be expected following such strong selection on permanent individual values in a relatively long-lived species (mean annual adult survival probability: ∼0.86, Frederiksen et al. 2008, giving an expected adult lifespan of ∼7 years) was not fully observed. Rather, mean permanent individual effects, and to some degree mean breeding values, decreased between the two ECEs, decreasing the expected degree of migration.

These decreases predominantly reflect effects of cohorts of new recruits that joined the focal adult population each year (Supporting Information S4). Since individuals recruiting following the ECEs hatched some years previously (median age of first reproduction: 3 years), they cause reversion towards mean breeding values of previous adults. This reversion could be accelerated by known episodes of reproductive selection against migrants (Grist et al. 2017; Acker et al. 2021b), and potentially also by as yet unquantified selection on pre-recruitment migration versus residence. Meanwhile, among-cohort variation in mean permanent individual effects on liability is likely to predominantly reflect lasting effects of developmental conditions rather than substantial shifts in non-additive genetic effects. These outcomes again point towards overarching effects of early-life conditions, not only in shaping emerging gene-by-environment interactions (outlined above), but also in shaping the phenotypic dynamics of seasonally-mobile populations in response to sequences of selective ECEs.

### Future prospects

High adult phenotypic repeatability of seasonal migration versus residence is increasingly observed in wild populations of diverse taxa (e.g. Kerr et al. 2009; Grist et al. 2014; Zúñiga et al. 2017; Sawyer et al. 2018; Lehnert et al. 2018). While such among-individual variation has not previously been explicitly partitioned into additive genetic versus different forms of permanent individual effect on liability- or phenotypic-scales, there is clearly scope for similar effects as uncovered in our study to act much more widely. The likely prominent role of permanent environmental effects should now drive new focus on early-life development of migration versus residence, encompassing advanced quantifications of multidimensional liability-scale reaction norms and their genetic basis and phenotypic-scale consequences. Such work can in future be facilitated by linking our conceptual framework and analytical methods with tracking technologies deployed on numerous related individuals, thereby integrating advances in movement ecology (Flack et al. 2022) with evolutionary quantitative genetics.

Meanwhile, our study highlights the limits of established quantitative genetic models that envisage current phenotypes as the sole substrate of selection on threshold traits (e.g. Roff 1998; de Villemereuil et al. 2016). By focusing solely on the fitness differential between the two phenotypes, such approaches filter out liability-scale covariance between relative fitness and breeding value that defines micro-evolutionary change. Fully predicting phenotypic responses in key threshold traits, such as migration versus residence, will therefore require quantifying selection on liability that would be hidden on the dichotomous phenotypic scale (Supporting Information S3: Figure S9). Such analyses could integrate fitness consequences of intrinsic phenotypic plasticity resulting from liability variation, including induced costs (DeWitt et al. 1998; Auld et al. 2010; Murren et al. 2015), alongside carry-over effects of previous phenotypic expression (Harrison et al. 2011), and indirect selection acting through other correlated traits (Lande and Arnold 1983). These ambitions are in principle achievable by extending our CRAM approach to jointly estimate environmental and additive genetic covariances between liability and fitness components, following principles of multivariate animal models (Morrissey et al. 2010, 2012; Stinchcombe et al. 2014). Our current assumptions of constant liability-scale genetic variance across seasonal occasions, environmental conditions and sexes can also be relaxed given more years of phenotypic and pedigree data, likely revealing further drivers and constraints on system dynamics. Yet, our current analyses provide the first insights into the fundamental quantitative genetic basis for joint plastic and evolutionary rescue of partially migratory populations experiencing changing circannual environments.

## Acknowledgements

We thank everyone who contributed to long-term field data collection, particularly Raymond Duncan, Sarah Fenn, Hannah Grist, Calum Scott, Jenny Sturgeon, and Moray Souter; and thank NatureScot for allowing work on the Isle of May National Nature Reserve. We thank Rita Fortuna and Thomas R. Haaland for useful comments on a manuscript draft. The current study was funded by Natural Environment Research Council (NERC; award NE/R016429/1 as part of the UK-SCaPE programme delivering National Capability), Norwegian Research Council (SFF-III grant 223257), NTNU and University of Aberdeen.

