## Supporting Information for "Components of micro-evolutionary and phenotypic change in seasonal migration versus residence in a wild population"

**Contents:**

**Part S1: Details of the study system and dataset (p. 2)**
- **S1.1 Phenotypic data collection (p. 2)**
- **S1.2 Preparation of the individual resighting histories (p. 4)**
- **S1.3 Structure of the pedigree data (p. 5)**

**Part S2: Details of the model (p. 10)**
- **S2.1 Full-annual-cycle multistate structure (p. 10)**
- **S2.2 Prior specifications (p. 12)**
- **S2.6 Additional model versions and checks (p. 13)**

**Part S3: Details of derived calculations (p. 17)**
- **S3.1 Decomposition of the phenotypic variance (p. 17)**
- **S3.2 Population-level dynamics of liabilities and resulting phenotypes (p. 20)**

**Part S4: Additional details of the results (p. 25)**
- **S4.1 General details of posterior sampling (p. 25)**
- **S4.2 Animal model part of the CRAM (p. 26)**
- **S4.3 Capture-recapture model part of the CRAM (p. 28)**

**Part S5: Literature cited (p. 36)**

### Part S1

### Details of the study system and dataset

#### S2.1 Phenotypic data collection

Winter-round geographic resightings allowed us to assign the phenotype migrant or resident when an individual was observed. The geographic range for resighting data collection (Figure S1) was defined based on historical dead recoveries and a prospective survey in winter 2008-09 which identified areas likely to cover winter locations of a large proportions of migrant shags that breed on the Isle of May ('IoM'; Grist et al. 2014, 2017; Acker et al. 2021a). Individuals have been observed from 540km north to 355km south of IoM at roosting sites on cliffs, rocky islets, and harbour walls. From winter 2009-10, observation effort during ~biweekly winter resighting surveys was directed at specific times, locations, and weather conditions, to maximise resighting efficiency. Extended details of the winter surveys are available in Grist et al. (2014, 2017) and Acker et al. (2021a,b, 2023) and appendices therein.

To control for spatio-temporal heterogeneity in observation failure due to variation in both observer effort and shags' observability, our model structure allowed resighting probability to vary through occasions and years across distinct states representing the residency area and five migrant areas (Figure S1; see also model details in Supporting Information S2). The residency area was defined as IoM and adjacent day roost sites (Figure S1; Acker et al. 2021a, 2023). The migratory areas encompass three regularly surveyed areas with typically high winter resighting effort, one geographically broad area representing all locations with low but non-zero resighting effort, and one 'ghost area' representing all migrant locations which presumably exist outside surveyed areas and hence where resighting probability is zero (e.g. Schaub et al. 2004; Figure S1).

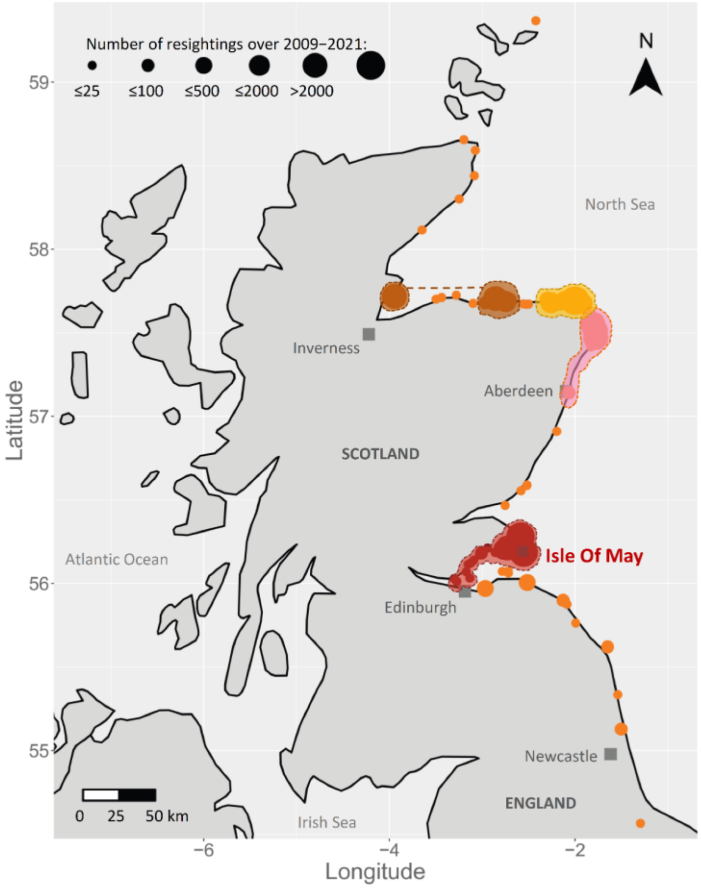


**Figure S1**. Locations of winter resightings of adult shags that bred on the Isle of May between 2009 and 2020 (modified from Acker et al. 2021a). Shaded zones show the four wintering areas that were surveyed ~biweekly, i.e. the residency area (red, IoM and nearby day-roosts) and three main migrant areas (brown, yellow, pink; data from the brown areas, linked by a dashed line, were pooled due to low sample sizes). Orange points show other locations of unstructured resightings pooled into one migrant area characterised by low expected resighting probability. The natural barrier formed by the Forth estuary (open water) separates the residency area from close south migratory locations on the southern shore of the Firth of Forth. Resident birds are typically resighted both on IoM and at coastal roosts close to their foraging locations along the north side of the Forth estuary. Individuals that were not resighted may have wintered in unobserved migrant locations which presumably exist on remote coasts, and are defined as "the ghost area".

Overall, the considerable resighting effort carried on throughout the winter led to relatively high resighting probability, ensuring collection of highly informative data to make probabilistic inference on among- and within-individual variation in migratory phenotype. Indeed, as shown in our previous analysis spanning 2009–2018, grand mean and range of resighting probability across all winter occasions and years was 0.47 (0.02–0.89) in the residency area, 0.66 (0.13-0.94) in regularly surveyed areas, and 0.34 (0.03–0.74) in the area pooling all irregularly surveyed locations (Acker et al., 2021a). We retrieved very similar estimates for 2009–2018 in the present analysis, and similar patterns for the three additional years analysed here 2018-2021 (see Supporting Information S4).

**S2.2 Preparation of the capture-recapture histories**

In our models, the annual cycle was divided into four winter occasions (1–4) plus the breeding season occasion ('B'). The exact date limits were 13$^{th}$ September to 14$^{th}$ October for occasion 1, 15$^{th}$ October to 14$^{th}$ November for occasion 2, 15$^{th}$ November to 31$^{st}$ December for occasion 3, and 1$^{st}$ January to 15$^{th}$ February for occasion 4, and 1$^{st}$ April to 30$^{th}$ June for occasion B. These time windows were carefully chosen to maximise the use of available phenotypic data while limiting violations of standard capture-recapture assumption of no movement and no mortality within occasion, and to enable our analyses to appropriately picture key aspects of within-winter variation in the expression of migration versus residence (see appendices of Acker et al. 2021a, 2022).

To compile individual resighting histories, all resightings of each focal individual in each occasion in each year were collapsed into a unique observation event (see full details of procedure in appendix of Acker et al. 2021a). Further, individual resighting histories were completed with information regarding observability of the focal individual. Indeed, individuals with worn or lost colour rings are unobservable from a distance in winter. Yet, summer conditions of observation allow observers to get closer to these individuals and read their virtually-unlosable metal ring and systematic effort is carried on to catch them and replace their colour ring. We considered these individuals (~3%) as unobservable in winter occasions up to the breeding season when their colour was replaced with a new one (see full details in appendices of Acker et al. 2021a, and archived model code).

##### S1.3 Structure of the pedigree data

We here provide additional description of the structure and content of the pedigree data.

*Founders, founders' descendants, and pedigree depth*

To estimate additive genetic variance, our analyses relied on a pedigree containing a large proportion of phenotype-informative individuals which were related to at least one other phenotype-informative individual (2058), alongside their additional ancestors (291).

More precisely, the utilised pedigree included 864 (37%) founders, of which 602 were phenotype-informative (26% of all individuals, 70% of the founders, 29% of the phenotype-informative individuals), and 262 were additional ancestors of phenotype-informative individuals (11% of all individuals, 30% of the founders, 90% of the additional ancestors). The 86 dummy parents (see main text) represented 4% of all individuals in the utilised pedigree, 10% of the founders, and 30% of the additional ancestors.

Further, among the 1485 remaining individuals comprising the founders' descendants (63% of the utilised pedigree), 1456 were phenotype-informative (62% of all individuals, 98% of the founders' descendants, 71% of the phenotype-informative individuals), and 29 were additional ancestors (1% of all individuals, 2% of the founders' descendants, 10% of the additional ancestors). Further among the founders' descendants, 515 individuals had both parents that were founders (22% of all individuals, 35% of the founders' descendants), and 660 had only one of their two parents that was a founder (28% of all individuals, 44% of the founders' descendants).

The utilised pedigree spanned four overlapping generations: 1207 individuals had no offspring in the pedigree (51% of all individuals), 848 (36%) had at least one offspring, 267 (11%) had at least one grand-offspring, and 27 (1%) had at least one great-grand-offspring.

*Parents, sibships, and broods*

Our utilised pedigree contained many parent-offspring links, and sibling links from the same or different broods. Such links are particularly useful for estimating additive genetic variance because they provide much information on the degree to which genetic relatedness is associated with phenotypic resemblance. More precisely, the utilised pedigree contained 789 unique parent pairs and 272 full sibships (see distribution of offspring number in Table S1). In total, 634 individuals had full siblings (27% of all individuals, 42% of the founders' descendants). In the founders' descendants, 975 individuals (66%) had at least one sibling (half or full) while 510 individuals (34%) had none.

**Table S1**. Distribution of the number of offspring in the utilised pedigree, across pairs (i.e. two identified parents), broods, females, and males.

| Number of offspring | Frequency | | | |
|---|---|---|---|---|
| | Pairs | Broods | Females | Males |
| 1 | 517 (65.5%) | 1601 (88.1%) | 326 (48.6%) | 306 (45.1%) |
| 2 | 204 (25.9%) | 191 (10.5%) | 199 (29.7%) | 191 (28.2%) |
| 3 | 53 (6.7%) | 25 (1.4%) | 89 (13.3%) | 93 (13.7%) |
| 4 | 11 (1.4%) | 0 | 37 (5.5%) | 55 (8.1%) |
| 5 | 2 (0.3%) | 0 | 14 (2.1%) | 20 (2.9%) |
| 6 | 1 (0.1%) | 0 | 1 (0.1%) | 8 (1.2%) |
| 7 | 1 (0.1%) | 0 | 3 (0.4%) | 1 (0.1%) |
| 8 | 0 | 0 | 0 | 1 (0.1%) |
| 9 | 0 | 0 | 1 (0.1%) | 1 (0.1%) |
| 10 | 0 | 0 | 0 | 2 (0.3%) |
| 11 | 0 | 0 | 0 | 0 |
| 12 | 0 | 0 | 1 (0.1%) | 0 |

The diversity of these links further provided us with relevant information to check whether shared developmental micro-environments associated with the brood, mother, or father identity also caused resemblance among close kins that could have inflated our estimates of additive genetic variance (Table S1; see also Supporting information S2).

More precisely, there were 216 broods with at least 2 individuals, representing a total of 457 individuals (31% of the founders' descendants) that had same-brood siblings (Table S1).

In the utilised pedigree, there were 693 mothers in total, and 345 maternal sibships (number of mothers with several offspring; see details in Table S1), including 173 maternal half-sibships (number of mothers that have several offspring with different males). These corresponded to a total of 931 individuals that had maternal siblings (40% of all individuals, 63% of the founders' descendants), including 297 that had maternal half-siblings (13% of all individuals, 20% of the founders' descendants).

Further, there were 678 fathers in total, and 372 paternal sibships (number of fathers with several offspring; see details in Table S1), including 252 paternal half-sibships (number of fathers that have several offspring with different females). These corresponded to a total of 1073 individuals that had paternal siblings (46% of all individuals, 72% of the founders' descendants), including 439 that had paternal half-siblings (19% of all individuals, 30% of the founders' descendants).

Moreover, there were 345 grand-mothers and 374 grand-fathers (15% and 16% of all individuals, respectively).

*Pedigree links among phenotype-informative individuals*

As expected given above-detailed elements of pedigree structure, most of the 8618 pairwise links among phenotype-informative individuals in the utilised pedigree (i.e. in the A matrix) corresponded to close additive genetic relatedness (Table S2, Figure S2).

**Table S2**. Distribution of the additive genetic relatedness values across all non-zero pairwise links between phenotype-informative individuals in the utilised pedigree.

| Additive genetic relatedness | Frequency of link | Proportion of all links |
|---|---|---|
| 0.00390625 | 8 | 0.09% |
| 0.0078125 | 51 | 0.6% |
| 0.015625 | 181 | 2.1% |
| 0.03125 | 418 | 4.9% |
| 0.0625 | 883 | 10.2% |
| 0.125 | 1704 | 19.8% |
| 0.1875 | 3 | 0.03% |
| 0.25 | 2681 | 31.1% |
| 0.375 | 1 | 0.01% |
| 0.5 | 2686 | 31.1% |
| 0.75 | 2 | 0.02% |

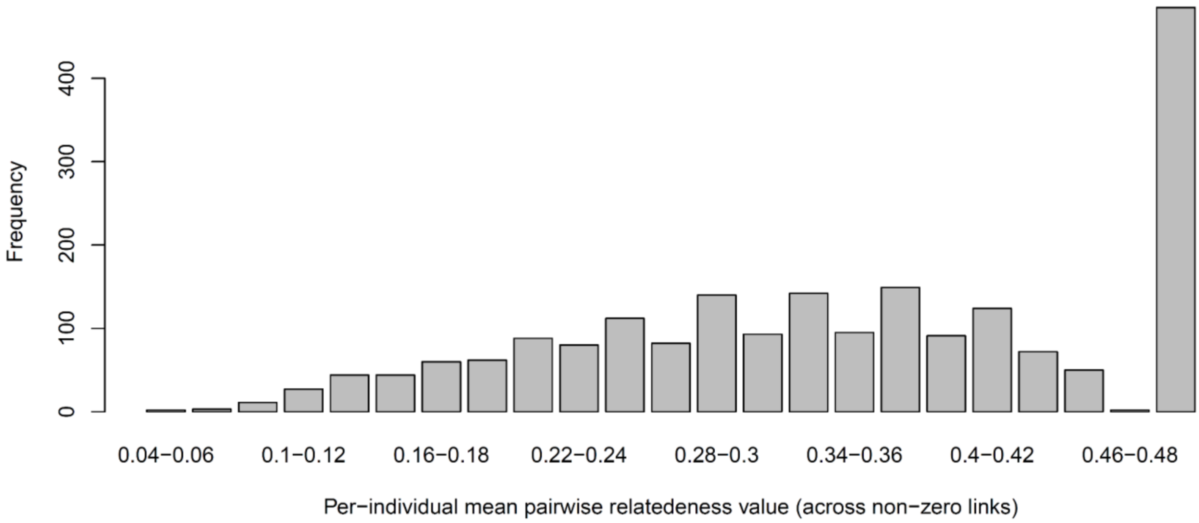


**Figure S2**. Distribution of the per-individual mean pairwise additive genetic relatedness values across non-zero links with other phenotype-informative individuals in the utilised pedigree.

The distribution of the per-individual number of pairwise links with other phenotype-informative individuals resembled an exponential distribution, with frequencies decreasing monotonically with the number of links (Figure S3).

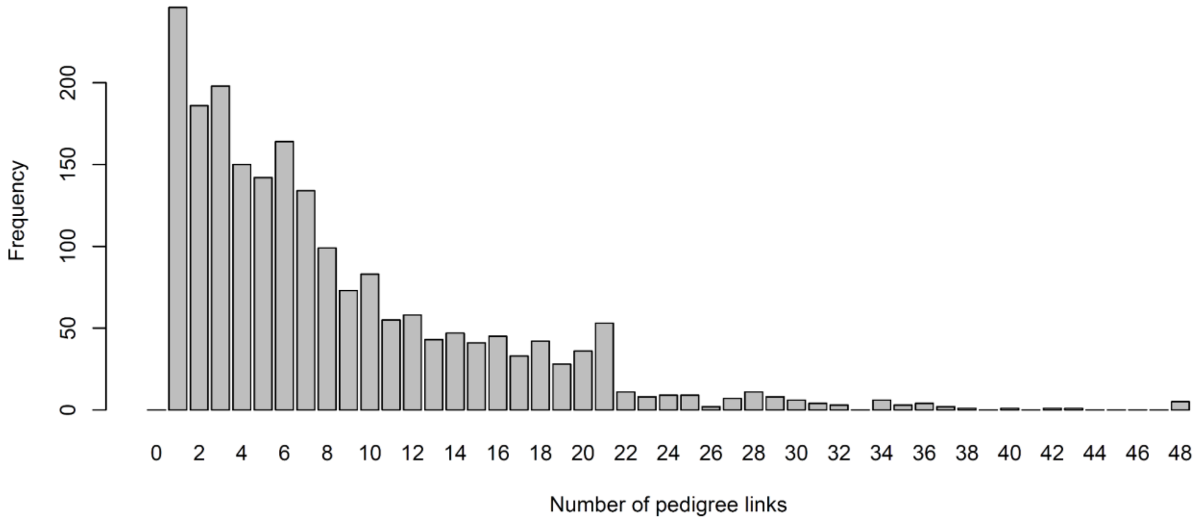


**Figure S3**. Distribution of the per-individual number of pairwise links between phenotype-informative individuals.

*Pedigree accuracy*

We used social parentage data, as commonly done in monogamous birds (e.g. Sheldon et al. 2003; Teplitsky et al. 2008; Charmantier et al. 2011; Husby et al. 2011), yet this might induce pedigree errors, due to non-exhaustiveness in individual identification and extra-pair paternity ('EPP').

However, we expected few pedigree gaps between phenotype-informative individuals in our study, given the very high proportion of known parents (see main text). Further, only 0–10% EPP were found in other Phalacrocoracidae species (Imperial shag, *Leucocarbo atriceps*: Calderon et al. 2012; great cormorants, *Phalacrocorax carbo*: Minias et al. 2015). These values match the ~0–20% EPP rates found by an early study on a subpart of the IoM population (Graves et al. 1993), but these estimates were based only on one minisatellite and would require to be updated with newer techniques and a larger dataset. Overall, as shown by simulation work, such low EPP rates are expected to cause minimal underestimation of additive genetic variance (Charmantier and Réale 2005; Firth et al. 2015).

### Part S2

### Details of the model and derived calculations

#### S2.1 Full-annual-cycle multistate structure

Our CRAM implies a full-annual-cycle structure of nested timescales (Figure S4).

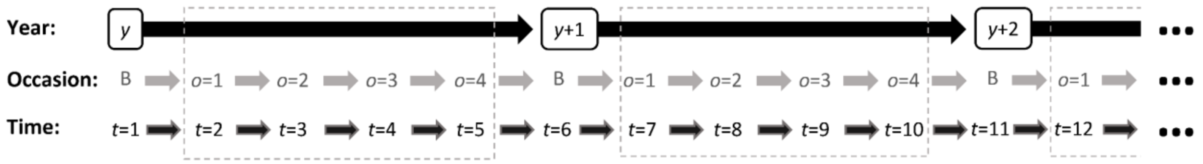


**Figure S4**. CRAM timeframes. Time (*t*) goes through breeding season (B) and winter occasions (*o*) across years (*y*). While each year in the capture-recapture model part of the CRAM includes B, the animal model part focuses on winter, i.e. when migration is expressed (dashed rectangle).

While the animal model part of the CRAM focuses on phenotypic variation in migration versus residence occurring in winter (Figure S4), the capture-recapture model part of the CRAM represents a full-annual-cycle process of transition between states (Figure S5). This allows to jointly model survival and movement throughout the years, and resulting spatio-temporally heterogeneous observations in the study area, as detailed by diagrams of fates in Figure S6.

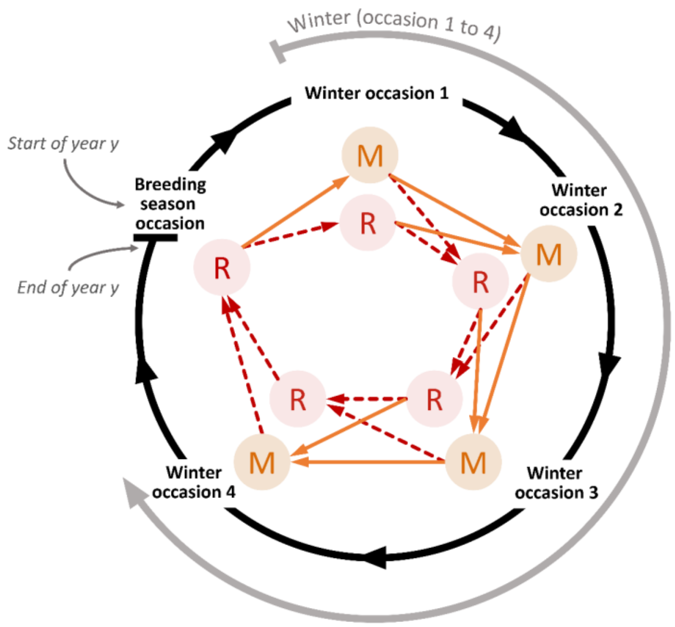


**Figure S5**. Full-annual-cycle structure of our CRAM. In the breeding season, all individuals are resident (R). Within the winter, surviving individuals can switch between R and migrant (M).

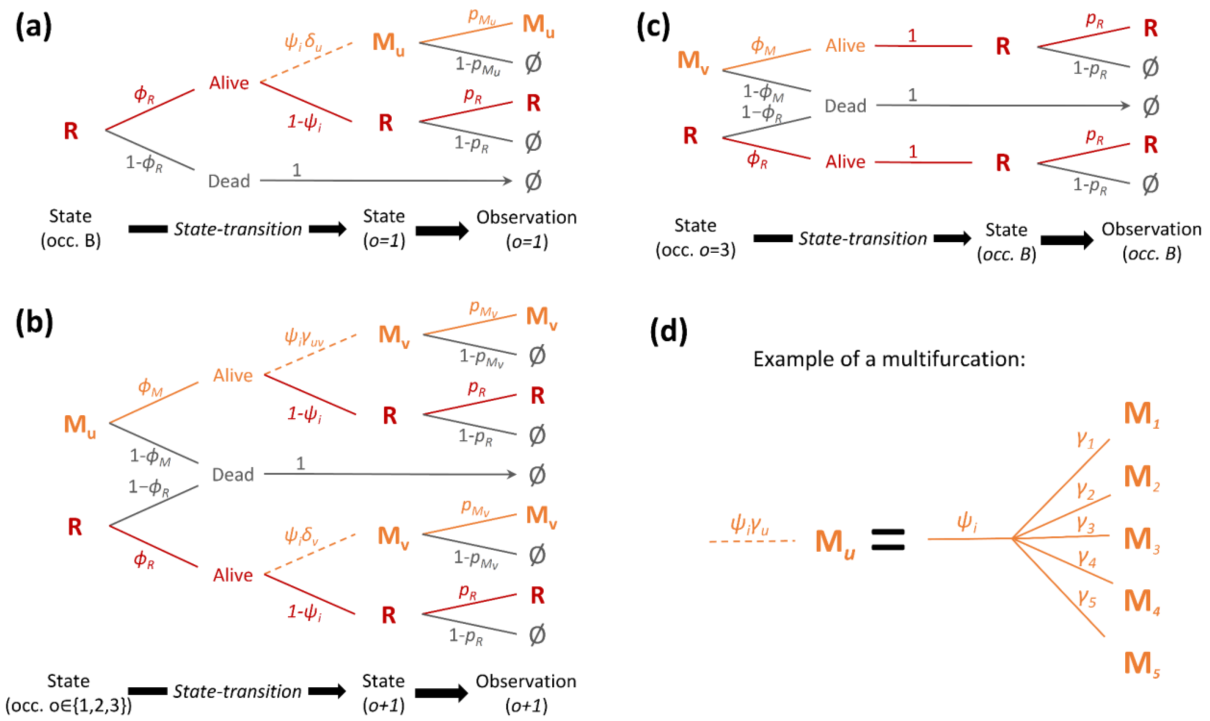


**Figure S6**. Diagram of fates for individual *i* throughout the full-annual-cycle in the CRAM. Panel (a) shows the possible transitions from the breeding season (B) to winter occasion 1, and subsequent observations. Here, all individuals start in the resident state (R) and can either remain resident in the first winter occasion or go to migrant area *u* ($M_u$; $u \in [\![1, 5]\!]$), or die (and hence end un in dead state, D. Arrows indicate possible paths in the state-transition and observation steps, with corresponding probabilities indicated as arrows' indices; dashed arrows symbolise multifurcations. Panel (b) shows the possible from winter occasion 1, 2, or 3 to the next winter occasion. Here, individuals can switch between resident and migrant areas (or remain in the same area; and accordingly, *u* can be either different from or equal to *v*). Panel (c) shows the possible transitions from winter occasion 4 to the following breeding season. Here, all individuals must go back to the residency area. Panel (d) shows an example of unfolded multifurcation. In all panels, parameters are elementary probabilities: $\phi_R$ and $\phi_M$ for survival of residents and migrants at *t*, $\varepsilon$ for departure (from the residency area), $\psi_i$ for expressing the migrant phenotype (note: this probability is specific to individual *i*), $\delta_u$ for moving to migratory area *u* ($u \in [\![1, 5]\!]$) conditional on departure ($\Sigma(\delta_u) = 1$), $\gamma_{uv}$ for switching from migratory area *u* to *v* ($v \in [\![1, 5]\!]$, *u* can be equal to *v*, and for $u \neq v$: $\gamma_{uv} = \frac{1-\gamma_{uu}}{4}$), $p_R$ and $p_{Mu}$ for resighting of residents and migrants (in area *u*; $p_u = 0$ if *u* is the 'ghost area'). These parameters can be occasion- and/or time-dependent (i.e. occasion × year-dependent; see main text).

**S2.2 Prior specifications**

We used uniform priors for all parameters in the capture-recapture part of the CRAM (Acker et al. 2021a). Here, all parameters were probabilities, either single probabilities of binary outcomes (e.g. survival probability of residents on a given time steps) or sets of probabilities of categorical outcomes, i.e. probability simplexes summing up to 1 (e.g. set of destination probabilities $\delta$ conditional on expressing the resident phenotype in a given winter occasion). Accordingly, we used uniform(0,1) distribution as the prior for any single probability of a binary outcome, and Dirichlet(1,…,1) as the prior for any set of probabilities of a categorical outcome. By using such priors, we remained objective regarding our general preconceptions on the values that the parameters could take (Acker et al. 2021a).

We used weakly informative priors for all parameters in the animal model part of the CRAM. Specifically, we used Student's $t$ distribution with 4 degrees of freedom, location 0 and scale 1, Student-t(4,0,1), as the prior for the sex×occasion-specific population-level intercepts on the standardized liability sale $\mu_{os}^{*}$. Such a prior is typically recommended for regression coefficients in logistic or probit regressions (Ghosh et al. 2018). During preliminary explorations, we also verified that our results were robust when we changed this prior to Normal(0,1) or Normal(0,2.5). For the additive genetic variance and permanent individual standard deviations, $\sigma_a^*$ and $\sigma_b^*$, we used the half-Student-t(4,0,1), i.e. Student-t(4,0,1) truncated to only have nonzero probability density for values greater than 0. Such a prior is typically recommended for variance parameters in hierarchical models (Gelman et al. 2006). During preliminary explorations, we also verified that variance estimates from our models were robust when we changed the scale of the half-Student-t distribution (to 0.75 and 1), or when we replaced the half-Student-T distribution to either half-Normal(0.075), half-Normal(0,1), half-Normal(0,1.5), half-Normal (0,2), half-Cauchy(0,1), half-Cauchy(0,5), exponential(1), or exponential(4.5), or Inverse-Gamma(0.001,0.001). Yet, the half-Student-t(4,0,1) provided the best performance in terms of Monte Carlo sampling. Notably, although the other priors led to similar posterior samples of $\sigma_a^{*2}$ and $\sigma_b^{*2}$, most of them implied Markov chains with divergent iterations (likely resulting in combination with relatively low power for estimating $\sigma_a^{*2}$, the key parameter on which CRAM inference is particularly data-demanding). The half-Student-t(4,0,1) did not lead to any divergent iterations.

#### S2.3 Additional model versions and checks

*Models with brood, maternal, or paternal effects*

To check whether common early-life environmental effects among close relatives could have inflated our estimate of $\sigma_a^{2*}$, we ran three additional models: one model which included brood effects, one model which included maternal effects, and one model which included paternal effects. For each of these models, the animal model part of the CRAM was formulated as:

$$probit(\psi_{io}) = \mu_{os_i}^* + a_i^* + b_i^* + k_{idx(i)}^*, \quad \text{(S1)}$$

where parameters are as in equation 11 of main text, $k$ is the focal kin effect (i.e. either brood, maternal, or paternal effect) and $idx(i)$ is the focal kin identity of individual $i$ (i.e. brood, mother, or father identity), and $k_{idx(i)}^* \sim \mathcal{N}(0, \sigma_k^2)$.

The posterior distributions of $\sigma_a^{2*}$ in each of these models and in the model presented in main text (which did not include any kin effect) were similar, and there was no or weak evidence of kin effects (Figure S7). Specifically, in all these models the additive genetic variance still clearly peaked away from zero, while the mode of the focal kin variance was always on zero (Figure 7), indicating that kin variance was negligible. The estimates of $\sigma_a^{2*}$ were 0.32 [0.01,0.73], 0.28 [0.00,0.68], and 0.27 [0.00,0.68] for the models with brood, maternal, and paternal effects, respectively (vs. 0.35 [0.01, 0.74] in the model with no kin effect; Table 1 of main text). The estimates of $\sigma_k^{2*}$ were 0.06 [0.00,0.28], 0.05 [0.00,0.20], and 0.09 [0.00,0.34], respectively, for the brood variance, maternal variance, and paternal variance. Note that here, MCMC sampling was particularly difficult, and there were divergent iterations in the Markov chains. This is likely because the CRAM inference on liability-scale variances of both additive genetic and kin effects is particularly data-demanding and the size of our dataset probably lies at the lower margin of what is necessary to estimate the components of interest.

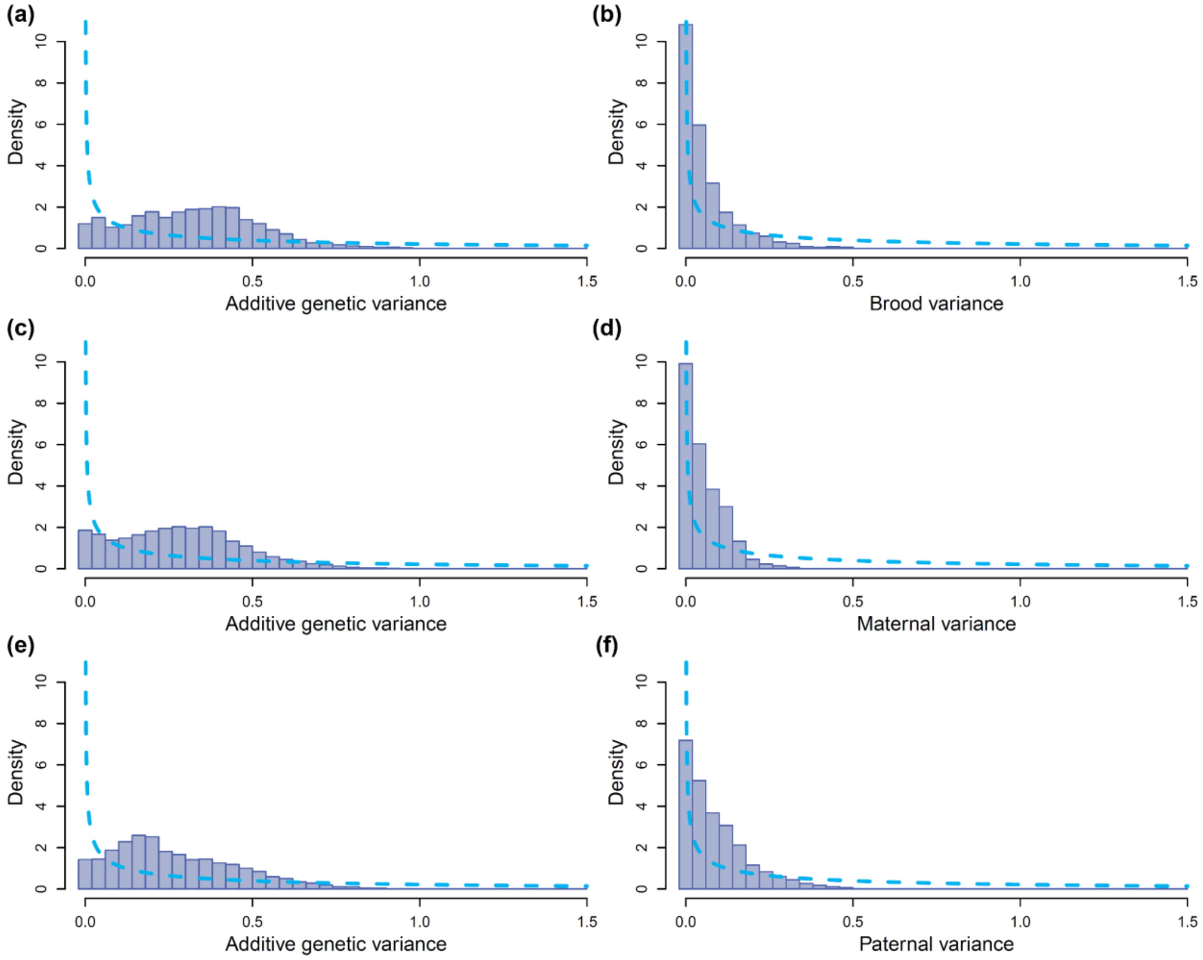


**Figure S7**. Prior (dotted line) and posterior (bars, MCMC samples) distributions of (a, c, e) additive genetic variance ($\sigma_a^{*2}$) and (b) brood, (d) maternal, (f) paternal variance ($\sigma_k^{*2}$) in liability for seasonal migration versus residence. Each of these three pairs of estimates ($\sigma_a^{*2}$ and $\sigma_k^{*2}$) come from each of the three additional models that included the focal kin effect. Estimates are on the standardised liability scale, where the unit is one standard deviation of the temporary residuals of the liability ($\sigma_\varepsilon$). These graphs were scaled to facilitate comparisons between the posterior distributions of $\sigma_a^{*2}$ and $\sigma_k^{*2}$ (and note that the present scales are substantially different from those in Figure 4 of main text).

*Control model with randomised A matrix*

Estimation of among-individual variance in binomial generalised linear models, including binomial animal models, is notoriously challenging. It is known from the animal breeding literature that relatively small datasets containing few individuals and/or few occasions of within-individual variation in phenotypic expression per individual may cause overestimation of total individual variance and additive genetic variance (Ødegård et al. 2010). The most pathological inferences leading to large bias and misleading uncertainty in the estimates may occur when the data on dichotomous phenotypic variation are strictly cross-sectional (i.e., only one instance of phenotypic expression per individual; Moreno et al. 1997; Ødegård et al. 2010). But estimations are expected to be satisfactory for datasets that contain many individuals and more than one effective phenotypic measurement per individual (Moreno et al. 1997). Nonetheless, it is reasonable to expect that pedigrees collected in wild populations can provide greater informativity per individual, thanks to greater connectiveness among individuals and diversity in relatedness values.

Here, given the large number of individuals and multiple repetitions of reversible trait expression throughout the years, we were confident that our dataset would provide enough information to robustly estimate total individual variance underlying the expression of migration vs. residence (e.g. Fay et al. 2022). And indeed, in previous analyses of fewer years of data, we already found strong evidence of individual phenotypic repeatability in migration versus residence within and between years, and distinctive structured patterns of among-individual variation in within-individual phenotypic variation that matched the threshold trait model (Acker et al. 2023). Further, simulations of a capture-recapture dataset using the same data structure, and estimates of survival and detection obtained in previous studies, demonstrated that our model was able to accurately estimate total individual variance alone (without pedigree data, i.e. following equation 11 of main text). Further using a highly informative simulated pedigree, the model was also able to accurately decompose this variance into additive genetic variance and permanent individual variance. Given the relatively high completeness of our pedigree data (see Wolak and Reid 2017 for comparisons with other wild population pedigrees), we were also confident that our real pedigree data would allow estimating additive genetic variance with little bias and satisfactory precision.

Nevertheless, to verify that our inference was not flawed by any systematic overestimation of the additive genetic variance on the standardised liability scale, $\sigma_a^{2*}$, we ran a 'control version' of our analysis, where we fitted our model to the real data after having randomly shuffled individual identities in the A matrix. Here, we retrieved an estimate of permanent individual variance that was similar to our estimate of total individual variance obtained with the real data (3.17 [2.71,3.69]; Figure S8). Further, the hence estimated additive genetic variance (0.12 [0.00,0.38]) was clearly negligible, as indicated by the posterior distribution which clearly peaked on zero (Figure S8). In contrast, the posterior distribution of the additive genetic variance obtained using the real pedigree data was clearly away from zero (Figure 4 of main text). In conclusion, the 'control version' of our analysis confirmed that our results obtained with the real data had not been spuriously generated.

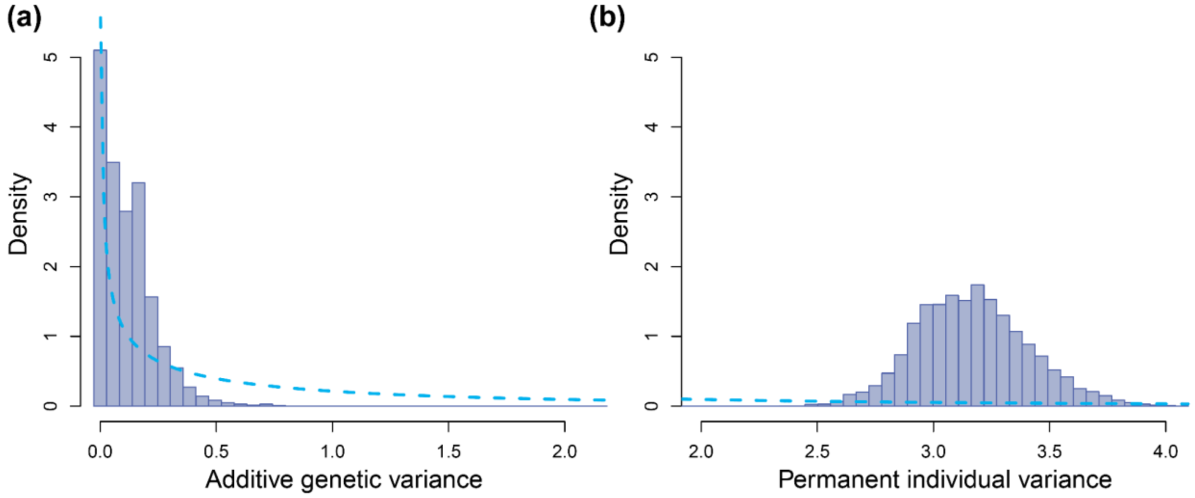


**Figure S8**. Prior (dotted line) and posterior (bars, MCMC samples) distributions of (a) additive genetic variance ($\sigma_a^{*2}$) and (b) permanent individual variance ($\sigma_b^{*2}$) in liability for seasonal migration versus residence, obtained from a 'control version' of the model run after having randomised individual identities in the A matrix. Estimates are on the standardised liability scale, where the unit is one standard deviation of the temporary residuals of the liability ($\sigma_\varepsilon$).

### Part S3

### Details of derived calculations

#### S3.1 Decomposition of the phenotypic variance

*Formulas for deriving phenotypic means and variances from liability-scale estimates*

Given the threshold-trait model, the expected population phenotypic mean $\bar{z}$ is the probability that the liability $\ell$ of an individual taken randomly in the population is above the threshold. In other words, this is the area above 0 under the probability density curve of a normal distribution with mean $\mu^*$ (overall population mean standardised liability) and standard deviation $\sigma_\ell^*$ (total trait variance on the standardised liability scale):

$$\bar{z} = 1 - F(0, \mu^*, \sigma_\ell^*) = F(\mu^*, 0, \sigma_\ell^*) = F\left(\frac{\mu^*}{\sigma_\ell^*}\right) \quad \text{(S2)}$$

where $F(0, \mu^*, \sigma_\ell^*)$ is the cumulative distribution function of a normal distribution with mean $\mu^*$ and standard deviation $\sigma_\ell^*$ evaluated at 0, and $F\left(\frac{\mu^*}{\sigma_\ell^*}\right)$ is the cumulative distribution of the standard normal distribution evaluated at $\frac{\mu^*}{\sigma_\ell^*}$.

By extension, the expected phenotype of an individual with a known effect *e* applied to its liability (with $e \sim \mathcal{N}(0, \sigma_e)$, taking value $e^* = e/\sigma_\varepsilon$ on the standardised liability scale), $E(z|e)$, is given by:

$$E(z|e) = F\left(\frac{\mu^* + e^*}{\sqrt{\sigma_\ell^{*2} - \sigma_e^{*2}}}\right). \quad \text{(S3)}$$

Here, it can be noted that if *e* is the total deviation applied to individual *i* at time *t* (i.e. if $e^* = a_i^* + b_i^* + \varepsilon_{it}^*$, and hence $\mu^* + e^* = \ell_{it}^*$), then equation S3 becomes the deterministic threshold model. Indeed, in this specific case $\sigma_e^* = \sqrt{\sigma_a^{*2} + \sigma_b^{*2} + 1} = \sigma_\ell^*$, which leads to $E(z|e) = E(z|\ell_{it}) = F(\infty * \ell_{it}^*)$, which is 0 if $\ell_{it}$<0 and 1 if $\ell_{it}$>0.

By definition (and see de Villemereuil et al. 2016), the variance of $E(z|e)$, i.e. $V_e$ the phenotypic variance due to liability-scale variation in effect $e$, is:

$$V_e = Var(\,E(z|e)) = \int (E(z|e) - \bar{z})^2 f_{\mathcal{N}}(e^*, 0, \sigma_e^*) de^*, \qquad \text{(S4)}$$

where $f_{\mathcal{N}}(e^*, 0, \sigma_e^*)$ is the probability density function of a normal distribution with mean 0 and standard deviation $\sigma_e^*$ evaluated at value $e^*$. It can be shown (see proof below) that this integral simplifies to:

$$V_e = \bar{z}(1 - \bar{z}) - 2T\left(\frac{\mu^*}{\sigma_\ell^*}, \frac{\sqrt{\sigma_\ell^{*2} - \sigma_e^{*2}}}{\sqrt{\sigma_\ell^{*2} + \sigma_e^{*2}}}\right), \qquad \text{(S5)}$$

where $T(h,c)$ is Owen's T function (Owen 1956; see below). Since $\forall h\ T(h,0) = 0$, we can directly note that, if $e = a_i + b_i + \varepsilon_{it}$ and hence for $\sigma_e^{*2} = \sigma_\ell^{*2}$, then we retrieve the well-known Bernoulli variance formula $V_z = \bar{z}(1 - \bar{z})$.

Equation S5 provides a simple general solution for estimating any phenotypic-scale variance resulting from any given liability-scale random effect or a linear combination of two or more liability-scale effects. Accordingly, equation S5 allowed us to directly calculate $V_a$, $V_b$, $V_\varepsilon$, $V_x$, and $V_z$ (given $\mu^*$, and $\sigma_a^*$, $\sigma_b^*$, $\sigma_\varepsilon^*$, $\sigma_x^*$, and $\sigma_\ell^*$). Moreover, when a linear combination of two or more liability scales effects are considered, the resulting phenotypic variance includes both independent and interaction effects on phenotypes resulting from the underlying liability-scale effects. Specifically, here $V_x = V_a + V_b + V_{a\times\mathrm{b}}$, accordingly we could derive $V_{a\times\mathrm{b}}$ as $V_x - V_a - V_b$. Similarly, we derived $V_{a\times\varepsilon}$ as $V_{a+\varepsilon} - V_a - V_\varepsilon$, $V_{b\times\varepsilon}$ as $V_{b+\varepsilon} - V_b - V_\varepsilon$, and $V_{a\times\mathrm{b}\times\varepsilon}$ as $V_z - V_x - V_\varepsilon - V_{a\times\varepsilon} - V_{b\times\varepsilon}$.

Following Robertson (1950), and de Villemereuil et al. (2016), the additive genetic variance on the phenotypic scale, $V_A$ is given by:

$$V_A = f_{\mathcal{N}}(0, \mu^*, \sigma_\ell^*)^2 \sigma_a^{*2}. \qquad \text{(S5)}$$

Since $V_a$ is the sum of variances arising from the independent and interaction effects on phenotypes of liability scale additive genetic effects, we can derive the variance of interaction effects $V_{NA}$ as $V_a - V_A$.

As noted in main text, since our model included occasion×sex-specific population-level intercepts on the liability scale ($\mu^*_{os}$), in our study all the above calculations and resulting derived estimates were occasion×sex-specific.

*Proof of equation S5*

Given equation S4, we can express $V_e$ as the sum of three integrals:

$$V_e = L + M + N \tag{S6}$$

$$L = \int \left(E(z|e)\right)^2 f_{\mathcal{N}}(e^*, 0, \sigma^*_e) de^* \tag{S7}$$

$$M = \int -2E(z|e)\bar{z} f_{\mathcal{N}}(e^*, 0, \sigma^*_e) de^* \tag{S8}$$

$$N = \int \bar{z}^2 f_{\mathcal{N}}(e^*, 0, \sigma^*_e) de^* \tag{S9}$$

In addition, following solutions for integrals of Gaussian functions provided by Owen (1980):

$$\int F(h + gx)^2 f_{\mathcal{N}}(x, 0{,}1) dx = F\left(\frac{h}{\sqrt{1+g^2}}\right) - 2T\left(\frac{h}{\sqrt{1+g^2}}, \frac{1}{\sqrt{1+2g^2}}\right). \tag{S10}$$

$$\int F(h + gx) f_{\mathcal{N}}(x, 0{,}1) dx = F\left(\frac{h}{\sqrt{1+g^2}}\right), \tag{S11}$$

Equation S7 is equivalent to:

$$L = \int \left(E(z|e)\right)^2 f_{\mathcal{N}}(e^*, 0, \sigma^*_e) de^*, \tag{S12}$$

$$= \frac{1}{\sigma^*_e} \int F\left(\frac{\mu^* + e^*}{\sqrt{\sigma^{*2}_\ell - \sigma^{*2}_e}}\right)^2 f_{\mathcal{N}}\left(\frac{e^*}{\sigma^*_e}, 0{,}1\right) de^*.$$

And given equation S10, posing $x = e/\sigma^*_e$, $h = \mu^*/\sqrt{\sigma^{*2}_\ell - \sigma^{*2}_e}$, $g = \sigma^*_e/\sqrt{\sigma^{*2}_\ell - \sigma^{*2}_e}$, we get:

$$L = F\left(\frac{\mu^*/\left(\sqrt{\sigma^{*2}_\ell - \sigma^{*2}_e}\right)}{\sqrt{1 + \sigma^{*2}_e/\left(\sigma^{*2}_\ell - \sigma^{*2}_e\right)}}\right) - 2T\left(\frac{\mu^*}{\sigma^*_\ell}, \frac{1}{\sqrt{1 + 2\frac{\sigma^{*2}_e}{\sigma^{*2}_\ell - \sigma^{*2}_e}}}\right), \tag{S13}$$

$$= F\left(\frac{\mu^*}{\sigma_\ell^*}\right) - 2T\left(\frac{\mu^*}{\sigma_\ell^*}, \frac{\sqrt{\sigma_\ell^{*2} - \sigma_e^{*2}}}{\sqrt{\sigma_\ell^{*2} + \sigma_e^{*2}}}\right),$$

$$= \bar{z} - 2T\left(\frac{\mu^*}{\sigma_\ell^*}, \frac{\sqrt{\sigma_\ell^{*2} - \sigma_e^{*2}}}{\sqrt{\sigma_\ell^{*2} + \sigma_e^{*2}}}\right).$$

Further, equation S8 is equivalent to:

$$M = -\frac{2\bar{z}}{\sigma_e^*}\int F\left(\frac{\mu^* + e^*}{\sqrt{\sigma_\ell^{*2} - \sigma_e^{*2}}}\right) f_{\mathcal{N}}\left(\frac{e^*}{\sigma_e^*}, 0{,}1\right) de^*. \quad \text{(S14)}$$

Given equation S11, posing again $x = e/\sigma_e^*$, $h = \mu^*/\sqrt{\sigma_\ell^{*2} - \sigma_e^{*2}}$, $\mathrm{g} = \sigma_e^*/\sqrt{\sigma_\ell^{*2} - \sigma_e^{*2}}$, we get:

$$M = -2\bar{z}\, F\left(\frac{\mu^*/\left(\sqrt{\sigma_\ell^{*2} - \sigma_e^{*2}}\right)}{\sqrt{1 + \sigma_e^{*2}/(\sigma_\ell^{*2} - \sigma_e^{*2})}}\right), \quad \text{(S15)}$$

$$= -2\bar{z}\, F\left(\frac{\mu^*}{\sigma_\ell^*}\right),$$

$$= -2\bar{z}^2.$$

And, trivially, equation S9 is equivalent to:

$$N = \bar{z}^2. \quad \text{(S16)}$$

Therefore, equations S6, S13, S15, and S16 give:

$$V_e = \bar{z} - \bar{z}^2 - 2T\left(\frac{\mu^*}{\sigma_\ell^*}, \frac{\sqrt{\sigma_\ell^{*2} - \sigma_e^{*2}}}{\sqrt{\sigma_\ell^{*2} + \sigma_e^{*2}}}\right), \quad \text{(S17)}$$

$$= \bar{z}(1 - \bar{z}) - 2T\left(\frac{\mu^*}{\sigma_\ell^*}, \frac{\sqrt{\sigma_\ell^{*2} - \sigma_e^{*2}}}{\sqrt{\sigma_\ell^{*2} + \sigma_e^{*2}}}\right).$$

#### S3.2 Population-level dynamics of liabilities and resulting phenotypes

*Calculation of population-level mean effects*

For each individual $i$, our model allowed us to estimate the breeding value $a_i^*$ and permanent individual effect $b_i^*$, and resulting individual×occasion-specific standardised liability expectations

$\eta^*_{io}$ and phenotypic expectation $\psi_{io}$. Yet, at each time $t$ across the study period, the population comprised different sets of individuals. Indeed, different individuals entered the population in different years during the breeding season (Table S3), and individuals may die at different times throughout the study period. However, while individuals are known to have been alive until their last resighting, there is uncertainty regarding whether and when they died after that (see Methods; Table S3). This uncertainty can be expressed given model parameters as the probability $\dagger_{it}$ that individual $i$ was dead at time $t$ (see details below).

Accordingly, the time-specific population-level mean breeding value $\bar{a}^*_t$, mean permanent individual effect $\bar{b}^*_t$, mean standardised liability expectation $\bar{\eta}^*_t$, and expected proportion of migrants $\bar{\psi}_t$ were calculated as weighted means:

$$\bar{a}^*_t = \frac{\sum_i w_{it} a^*_i}{\sum_i w_{it}}, \bar{b}^*_t = \frac{\sum_i w_{it} b^*_i}{\sum_i w_{it}}, \bar{\eta}^*_t = \frac{\sum_i w_{it} \eta^*_{io(t)}}{\sum_i w_{it}}, \text{and } \bar{\psi}_t = \frac{\sum_i w_{it} \psi_{io(t)}}{\sum_i w_{it}}, \quad \text{(S18)}$$

where $w_{it}$ is the weight of individual $i$ at $t$ is 0 before the individual's entry in the dataset, 1 from its entry in the dataset to its last resighting, and 1-$\dagger_{it}$ after its last resighting; and $o(t)$ is the within-winter occasion (1, 2, 3, or 4) of to the focal time step.

**Table S3**. Annual numbers of individuals in the breeding season which are either (a) entering the dataset ("entries"), (b) already entered and known to be still alive because they have been resighted afterwards, or (c) already entered but possibly dead because they have never been resighted afterwards.

| | Year | | | | | | | | | | | | |
|---|---|---|---|---|---|---|---|---|---|---|---|---|---|
| | 2009 | 2010 | 2011 | 2012 | 2013 | 2014 | 2015 | 2016 | 2017 | 2018 | 2019 | 2020 | 2021 |
| (a) Entries | 721 | 159 | 263 | 302 | 145 | 165 | 113 | 112 | 190 | 159 | 171 | 76 | 0 |
| (b) Known alive | NA | 680 | 786 | 956 | 559 | 538 | 648 | 680 | 722 | 697 | 717 | 783 | 771 |
| (c) Possibly dead | NA | 41 | 94 | 187 | 886 | 1052 | 1107 | 1188 | 1258 | 1473 | 1612 | 1717 | 1805 |

Note that $\dagger_{it}$ increases with time from the last resighting, i.e. the longer the time since last resighting, the more likely an individual is to be dead. In our data, since resighting probability in the breeding season is typically very high (mean ~0.95; see Methods and Supporting Information S4), individuals that are not resighted in a given breeding season are highly likely to be dead, and individuals that are not seen two years in a row are almost certainly dead. Yet, there is greater uncertainty regarding exact timing of death within a given winter.

*Calculation of* $\dagger_{it}$

The probability $\dagger_{it}$ that an individual $i$ was dead (state '*D*') on time $t$ after its last resighting is a complex function of its phenotypic expectation $\psi_i$, its last observed location, and subsequent phenotype-dependent survival probabilities ($\phi$), migrant-area destination and movement probabilities ($\delta$ and $\gamma$), and resighting probabilities in the different areas ($p$). There is no simple general analytical formula for computing this complex individual×occasion×year probability, but it can be done using the well-known 'forward algorithm' for computing probabilities of hidden state variables in Markov models given a sequence of observation (Baum 1972; Juang & Rabiner 1986; Rabiner 1990; and see archived model code).

In general, this algorithm allows us to compute the probability that the state $S_{it}$ of individual $i$ at a given time $t$ (after, on, or before last sighting) was $k$, given its capture-resighting history $H_{i,1:t}$ from time 1 to $t$, *i.e.* $P(S_{it} = k | H_{i,1:t})$, as a function of probabilities of state transition between successive time steps (defined by $\psi_i$, $\phi$, $\delta$, and $\gamma$) and state-dependent observation ($p$). We can hence express the probability of an individual's entire capture-recapture history (i.e. $P(H_{i,1:E}) = \sum_k P(S_{iE} = k | H_{i,1:E})$, where $E$ is the end time point in the model), and hence the full likelihood of the model (i.e. $\prod_i \sum_k P(S_{iE} = k | H_{i,1:E})$).

This algorithm is also key in the calculation of $\dagger_{it}$ (for whichever $t$ after the last sighting of $i$). Indeed, according to the definition of conditional probability:

$$\dagger_{it} = P(S_{it} = \text{'}D\text{'} | H_{i,1:E}) = \frac{P\left(S_{it} = \text{'}D\text{'} \cap H_{i,1:E}\right)}{P\left(H_{i,1:E}\right)}. \quad \text{(S19)}$$

Moreover, given first-order Markovian dependence in an individual's state sequence:

$$\dagger_{it} = \frac{P\left(S_{it} = \text{'}D\text{'} \cap H_{i,1:(t-1)}\right) P\left(S_{it} = \text{'}D\text{'} \cap H_{i,t:E}\right)}{P\left(H_{i,1:E}\right)}. \quad \text{(S20)}$$

Yet, since here the last sighting of $i$ occurred before $t$, then by definition $P\left(S_{it} = \text{‘}D\text{’} \cap H_{i,t:E}\right) = 1$ (because $i$ was not resighted from $t$ to $E$, and dead individuals cannot be resighted). Following the state transition process defined in our model, we can then reformulate equation S20 as:

$$\dagger_{it} = \frac{\sum_k \left(P(S_{i(t-1)} = k \mid H_{i,1:(t-1)}) P(S_{it} = \text{‘}D\text{’} \mid S_{i(t-1)} = k)\right)}{P\left(H_{i,1:E}\right)}, \qquad \text{(S20)}$$

which can be computed given the model parameters with help of the forward algorithm (see archived model code for the practical application to our analysis).

*Implications of model assumptions on estimated changes in population-level mean effects*

Because $\dagger_{it}$ is calculated as a function of phenotype-dependent survival probability, it partly relies on the model assumption that phenotypes are the sole causation of survival selection. Indeed, when an individual is not resighted at or after $t$, the expectation that it survived from $t-1$ to $t$ will be dependent on its observed or expected phenotype at $t-1$ and the cross-sectional phenotype-specific estimate of survival probability between $t$ and $t-1$. However, the impact of this assumption is mitigated by the informativity of the full individual resighting history, implying that $\dagger_{it}$ should be closer to the true probability that individual $i$ was dead at time $t$ than what would have been expected solely from its phenotypic expectation and the phenotype-dependent survival probability. Moreover, until their last resighting, individuals are directly identified as survivors, and hence their weight in population-level trait dynamics is directly observed and does not imply any assumption on selection.

Nonetheless, the model assumption that phenotypes are the sole cause of survival selection likely led to some underestimation of selection that acted on the liability-scale. Indeed, here we estimated a survival probability of ~0.47 in resident and ~0.77 in migrants in late winter 2012-13 (Acker et al. 2021a; Supporting Information S4). Assuming that the late-winter population distribution of liabilities was a Normal with mean -0.2 and variance 4.3 (as found here: Table 1 of main text), we get a gradient of selection on liability of 0.09 (Figure S9). However, in fact we also previously found evidence indicating that underlying selection on the liability scale was likely stronger than would have been expected based solely on the fitness differential between the two phenotypes (see main text; Acker et al. 2023). We can illustrate this point by now assuming that, in late winter 2012-13, survival was a function of liability such that $logit(\phi) = 0.6 + 0.43\ell$, which results in the same phenotypic selection (survival probability is 0.47 in residents and 0.77 in migrants) but yields a gradient of selection on liability of 0.14 (Figure S9).

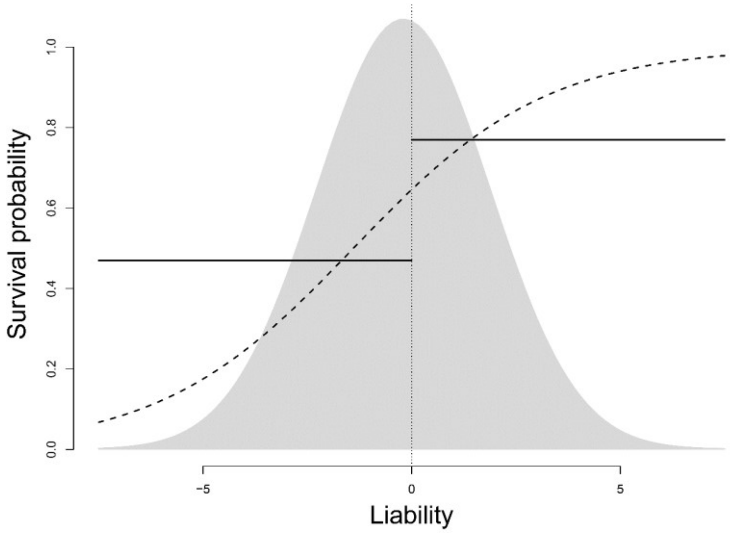


**Figure S9**. Discrepancy between liability-scale fitness as inferred from the survival differential between the two phenotypes (plain line; values are estimates for late winter 2012-13 in our study) vs. a hypothetical underlying continuous survival function on the liability scale (dashed line). The dotted vertical line shows the threshold. The background Gaussian density plot shows the expected distribution of liabilities in late winter in the population.

*Measuring the evidence for changes in $\bar{a}^*$ or $\bar{b}^*$*

We calculated temporal changes Δ in $\bar{a}^*$ and $\bar{b}^*$ between any two successive time points as:

$$\Delta(\bar{a}, t) = \bar{a}_t^* - \bar{a}_{t-1}^* \text{ and } \Delta(\bar{b}, t) = \bar{b}_t^* - \bar{b}_{t-1}^*.$$

In practice, as for any other derived parameters or quantities, Δ is calculated at each iteration of the MCMC chains (i.e. in each sample of the posterior distributions of the model parameters). This results in a full set of samples of the posterior distribution of the derived parameters. Here, to evaluate the evidence for a given direction of change in $\bar{a}^*$ or $\bar{b}^*$, we calculated the proportion of posterior samples of Δ which had the same sign as the posterior mean of Δ (i.e. we estimated the posterior probability that Δ had the same sign as its posterior mean).

### Part S4
### Additional details of the results

#### S4.1 General details of posterior sampling

Here, we present additional graphical and numerical summaries of the posterior distribution for parameters and derived quantities that were not shown in the main text. Detailed summaries of the posterior distributions of all parameters and derived quantities will be made available alongside complete posterior samples as 'Rdata' files (R Development Core Team, 2020), archived in Zenodo (see data accessibility statement in main text).

Posterior sampling was performed using 10 Markov chains with 1000 warmup iterations followed by 3000 monitored iterations (without thinning), yielding 30000 samples in total. No divergent iterations occurred during sampling of the main model presented in the manuscript. Other versions of the model run during preliminary exploratory analyses with different parameterisations and specifications suffered from divergent iterations, which were clearly associated with difficulties in estimating the additive genetic variance $\sigma_a^{*2}$. Here, the Gelman-Rubin-Brooks *r-hat* statistics (Brooks and Gelman, 1998) indicated good convergence of the chains, with all *r-hat* <1.01, except $\sigma_a^{*2}$ for which *r-hat* was ~1.02. Monte Carlo standard errors were all <10% of the standard deviation of the corresponding posterior sample. Effective sample sizes of the model parameters were all >10% of the total sample size, except for $\sigma_a^{*2}$ and the permanent individual variance $\sigma_e^{*2}$ for which effective sample sizes were 366 and 882, respectively. Nonetheless, effective sample sizes were much larger for all other parameters, and all >7500 (i.e. >25% of total sample size). Overall, these diagnostics indicated a satisfactory sampling that allowed reliable inference, although unsurprisingly they also pointed towards difficulties in the estimation of $\sigma_a^{*2}$ and $\sigma_e^{*2}$. To conclude, the uncertainty of the estimates due to imperfect (pseudorandom) sampling was negligible, including for $\sigma_a^{*2}$ and $\sigma_e^{*2}$. Precision of the sampling was sufficiently high so that all posterior means of the model parameters could be reported with at least 2 decimal place precision. See Lunn et al. (2012) and Gelman et al. (2013) for further information on MCMC methods for posterior sampling (and more generally on Bayesian data analysis).

#### S4.2 Animal model part of the CRAM

*Phenotypic variances*

In the main text, we provided graphical summaries for the variance decomposition on the phenotypic scale (Figure 5). Hereafter, we provide the corresponding numerical summaries for each component of phenotypic variance (Table S4).

**Table S3**. Estimates of the phenotypic variance components and associated proportion of total phenotypic variance (mean and 95% credible interval).

| Variance component | Sex | Occasion 1 | | Occasion 2,3,4 | |
|---|---|---|---|---|---|
| | | Absolute value | Proportion of $V_z$ | Absolute value | Proportion of $V_z$ |
| $V_z$ | F | 0.23 [0.22, 0.24] | 1 | 0.25 [0.25, 0.25] | 1 |
| | M | 0.19 [0.18, 0.21] | 1 | 0.25 [0.25, 0.25] | 1 |
| $V_a$ | F | 0.01 [0.00, 0.02] | 0.05 [0.00, 0.10] | 0.01 [0.00, 0.03] | 0.05 [0.00, 0.11] |
| | M | 0.01 [0.00, 0.02] | 0.05 [0.00, 0.09] | 0.01 [0.00, 0.03] | 0.05 [0.00, 0.11] |
| $V_b$ | F | 0.11 [0.09, 0.13] | 0.47 [0.40, 0.55] | 0.12 [0.10, 0.14] | 0.48 [0.41, 0.56] |
| | M | 0.09 [0.07, 0.11] | 0.46 [0.38, 0.54] | 0.12 [0.10, 0.14] | 0.48 [0.41, 0.56] |
| $V_{a\times b}$ | F | 0.01 [0.00, 0.01] | 0.03 [0.00, 0.05] | 0.01 [0.00, 0.01] | 0.02 [0.00, 0.04] |
| | M | 0.01 [0.00, 0.01] | 0.03 [0.00, 0.06] | 0.01 [0.00, 0.01] | 0.02 [0.00, 0.04] |
| $V_\varepsilon$ | F | 0.03 [0.03, 0.04] | 0.14 [0.13, 0.16] | 0.04 [0.03, 0.04] | 0.15 [0.13, 0.17] |
| | M | 0.03 [0.02, 0.03] | 0.13 [0.12, 0.15] | 0.04 [0.03, 0.04] | 0.15 [0.13, 0.17] |
| $V_{a\times\varepsilon}$ | F | 0.00 [0.00, 0.00] | 0.00 [0.00, 0.01] | 0.00 [0.00, 0.00] | 0.00 [0.00, 0.01] |
| | M | 0.00 [0.00, 0.00] | 0.01 [0.00, 0.01] | 0.00 [0.00, 0.00] | 0.00 [0.00, 0.01] |
| $V_{b\times\varepsilon}$ | F | 0.03 [0.02, 0.06] | 0.13 [0.07, 0.26] | 0.03 [0.02, 0.06] | 0.12 [0.07, 0.25] |
| | M | 0.03 [0.02, 0.06] | 0.15 [0.09, 0.29] | 0.03 [0.02, 0.06] | 0.12 [0.07, 0.25] |
| $V_{a\times b\times\varepsilon}$ | F | 0.04 [0.01, 0.05] | 0.17 [0.04, 0.22] | 0.04 [0.01, 0.06] | 0.17 [0.04, 0.22] |
| | M | 0.03 [0.01, 0.05] | 0.18 [0.04, 0.23] | 0.04 [0.01, 0.06] | 0.17 [0.04, 0.22] |
| $V_x$ | F | 0.13 [0.12, 0.13] | 0.55 [0.52, 0.57] | 0.14 [0.13, 0.14] | 0.56 [0.53, 0.58] |
| | M | 0.10 [0.09, 0.11] | 0.54 [0.51, 0.56] | 0.14 [0.13, 0.14] | 0.56 [0.53, 0.58] |
| $V_A$ | F | 0.01 [0.00, 0.02] | 0.05 [0.00, 0.10] | 0.01 [0.00, 0.03] | 0.05 [0.00, 0.11] |
| | M | 0.01 [0.00, 0.02] | 0.05 [0.00, 0.09] | 0.01 [0.00, 0.03] | 0.05 [0.00, 0.11] |

Notes: $V_z$ is the total phenotypic variance. $V_a$, $V_b$ and $V_\varepsilon$ are the phenotypic variances arising independently from liability-scale additive genetic effects $a$, permanent individual effects $b$, and temporary residual effects $\varepsilon$, respectively. $V_{a\times b}$, $V_{a\times\varepsilon}$, $V_{b\times\varepsilon}$, and $V_{a\times b\times\varepsilon}$ are the phenotypic variances arising from interactions between these liability-scale effects. $V_x$ is the total phenotypic-scale individual variance. $V_A$ is the phenotypic-scale additive genetic variance. In the sex column, 'F' is female, and 'M' is male.

*Temporal change in population mean liability-scale effects*

Here we provide the posterior probabilities $P_{\Delta}$ that the difference $\Delta$ $\bar{a}^{*}$ or $\bar{b}^{*}$ in between two consecutive time points had the same sign as its posterior mean (Table S5). We also provide a graphical summary of the values of $\bar{a}^{*}$ or $\bar{b}^{*}$ on the first winter occasion of each year in the subset of individuals that just entered the dataset and those that were already present in the dataset (Figure S10).

**Table S5**. Summary of the posterior probabilities $P_{\Delta}$ measuring the support for the evidence of a signed difference $\Delta$ in $\bar{a}^{*}$ or $\bar{b}^{*}$ between any two consecutive winter occasions throughout study period.

| Year | Occasion | Parameter | | Year | Occasion | Parameter | |
|---|---|---|---|---|---|---|---|
| | | $\bar{a}^{*}$ | $\bar{b}^{*}$ | | | $\bar{a}^{*}$ | $\bar{b}^{*}$ |
| 2009-10 | 1 | 0.66 | 0.87 | 2015-16 | 1 | 0.53 | 0.86 |
| | 2 | 0.60 | 0.78 | | 2 | 0.68 | 0.89 |
| | 3 | 0.53 | 0.66 | | 3 | 0.74 | 1.00 |
| | 4 | 0.53 | 1.00 | | 4 | 0.70 | 0.98 |
| 2010-11 | 1 | 0.76 | 0.92 | 2016-17 | 1 | 0.62 | 0.90 |
| | 2 | 0.61 | 0.67 | | 2 | 0.54 | 0.50 |
| | 3 | 0.75 | 0.86 | | 3 | 0.57 | 0.80 |
| | 4 | 0.60 | 1.00 | | 4 | 0.58 | 0.99 |
| 2011-12 | 1 | 0.58 | 0.77 | 2017-18 | 1 | 0.75 | 0.93 |
| | 2 | 0.61 | 0.62 | | 2 | 0.50 | 0.59 |
| | 3 | 0.51 | 0.62 | | 3 | 0.51 | 0.58 |
| | 4 | 0.56 | 1.00 | | **4** | **0.92** | **1.00** |
| 2012-13 | 1 | 0.59 | 0.68 | 2018-19 | 1 | 0.63 | 0.81 |
| | 2 | 0.71 | 0.77 | | 2 | 0.60 | 0.69 |
| | 3 | 0.65 | 0.82 | | 3 | 0.67 | 0.79 |
| | **4** | **0.92** | **1.00** | | 4 | 0.53 | 0.60 |
| | 1 | 0.64 | 0.89 | | 1 | 0.66 | 0.71 |
| 2013-14 | 2 | 0.57 | 0.80 | 2019-20 | 2 | 0.58 | 0.65 |
| | 3 | 0.54 | 0.60 | | 3 | 0.54 | 0.64 |
| | 4 | 0.65 | 0.73 | | 4 | 0.67 | 1.00 |
| 2014-15 | 1 | 0.74 | 0.96 | 2020-21 | 1 | 0.75 | 1.00 |
| | 2 | 0.67 | 0.70 | | 2 | 0.79 | 0.92 |
| | 3 | 0.62 | 0.57 | | 3 | 0.50 | 0.72 |
| | 4 | 0.66 | 0.98 | | 4 | – | – |

Note: Values for occasions spanning an episode of strong survival selection are shown in bold.

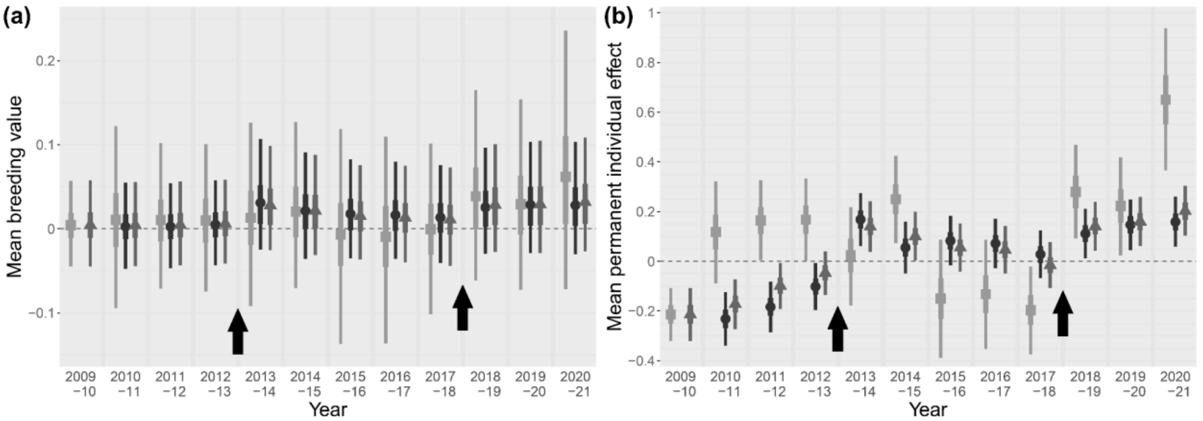


**Figure 6**. Estimated values of means of (a) breeding values ($\bar{a}$) and (b) permanent individual effects ($\bar{b}$) in the first winter occasion of each year among individuals that just entered the dataset in the previous breeding season (light grey squares) and those that were already in the dataset (dark grey circles). The third point in each year (grey triangles) shows the estimate for the full population (i.e. both subsets together, as shown on Figure 6 of the main text). Point estimates are posterior means, inner and outer line segments indicate 50% and 95% credible intervals. Black arrows point at the two episodes of strong survival selection that occurred in late-winter 2012-13 and 2017-18.

Here, we can see that the population-level mean essentially reflects the mean of the subset of composed individuals that were already present in the dataset in the previous year. Nonetheless, the accumulation of small changes due to the entry of new individuals do contribute to the long-term dynamics.

##### S4.3 Capture-recapture model part of the CRAM

*Survival probabilities*

Hereafter we provide graphical summaries of the posterior distribution of the probability of survival in residents and migrants (fig. S7 and S8, respectively). Comparisons of these two graphs clearly show the two events of high mortality that resulted in strong selection against residence in late-winter 2012-13 and 2017-18 (Acker et al. 2021a).

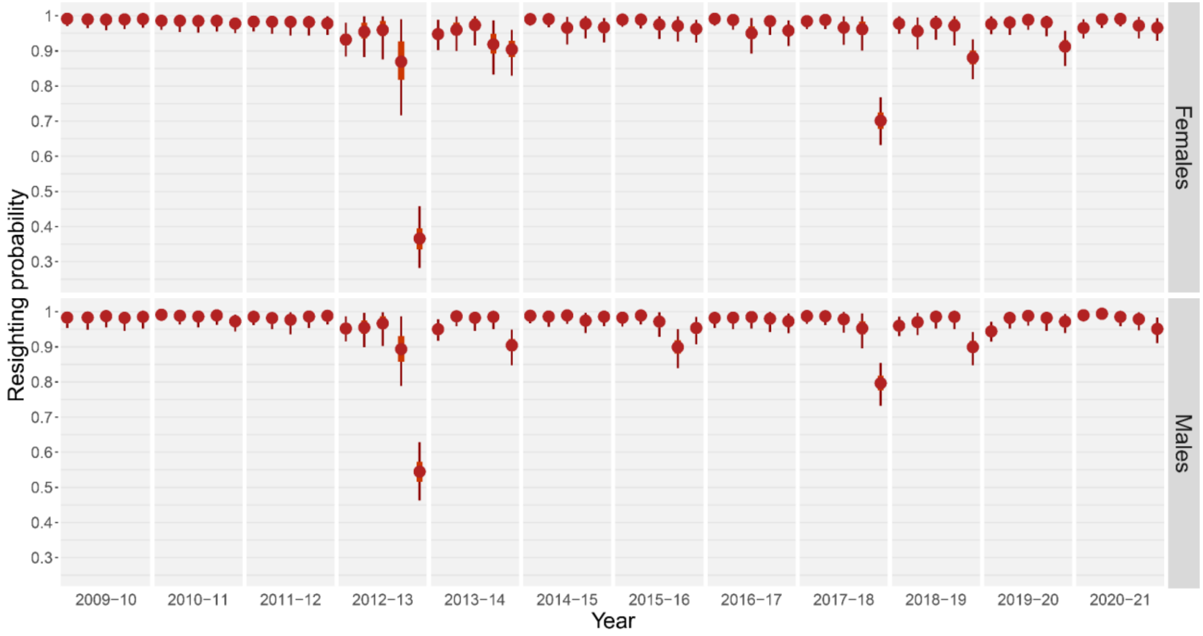


**Figure S7**. Probability of survival from one occasion to the next in female and male residents. In each year, the five successive points correspond to the successive occasions: the breeding season followed by the four winter occasions. Point estimates are posterior means, inner and outer line segments indicate 50% and 95% credible intervals.

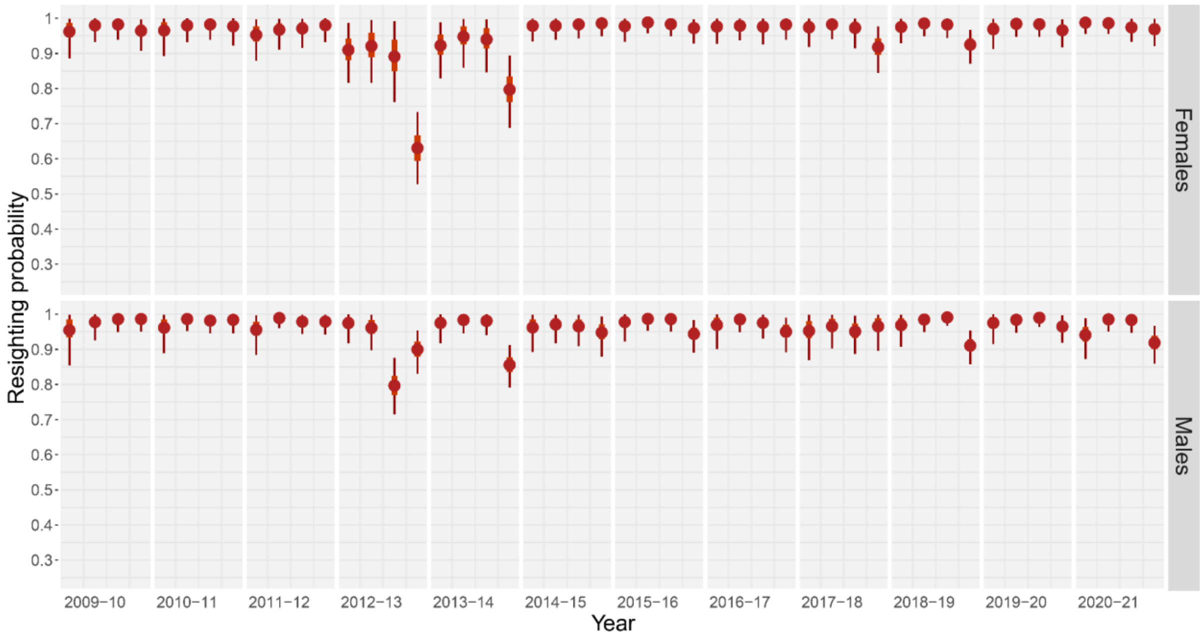


**Figure S8**. Probability of survival from one occasion to the next in female and male migrants. In each year, the four successive points correspond to the four winter occasions during which the migrant phenotype may be expressed by individuals. Point estimates are posterior means, inner and outer line segments indicate 50% and 95% credible intervals.

*Movement probabilities*

Hereafter we provide graphical summaries of the posterior distribution of the probability of moving in each migratory area (conditional on departure; fig. S9 to S13) and the probability of switching to another migratory area (conditional on not returning to the residency area; fig. S14). Remind that the probability of departing from the residency area or coming back from a migratory area are defined as the expression of the trait, which results from variation in liability specified by the animal model part of the CRAM.

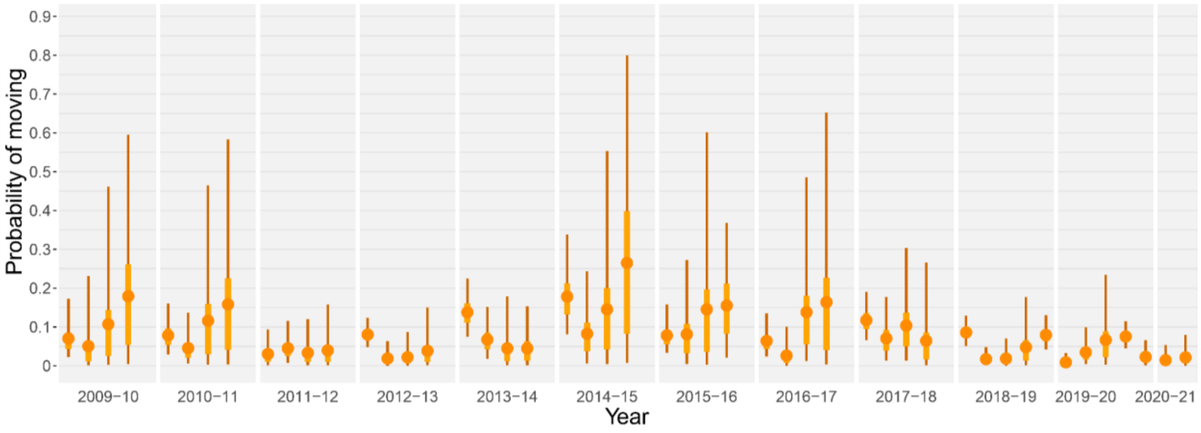


**Figure S9**. Probability of moving to migratory area 1 (oranges points on fig. S1), conditional on departure. This probability is constant across sexes. In each year, the four successive points correspond to the four successive winter occasions. Point estimates are posterior means, inner and outer line segments indicate 50% and 95% credible intervals.

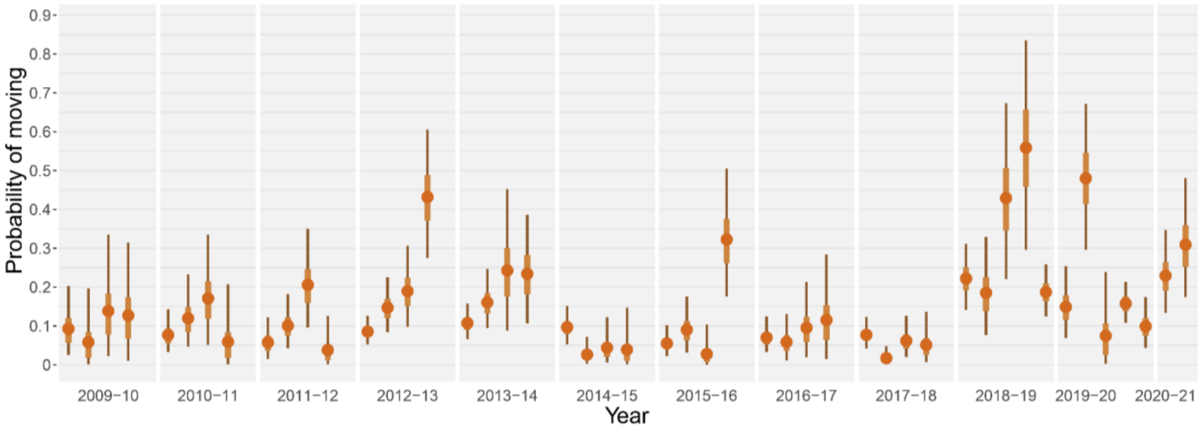


**Figure S10**. Probability of moving to migratory area 2 (brown area on fig. S1), conditional on departure. This probability is constant across sexes. In each year, the four successive points correspond to the four successive winter occasions. Point estimates are posterior means, inner and outer line segments indicate 50% and 95% credible intervals.

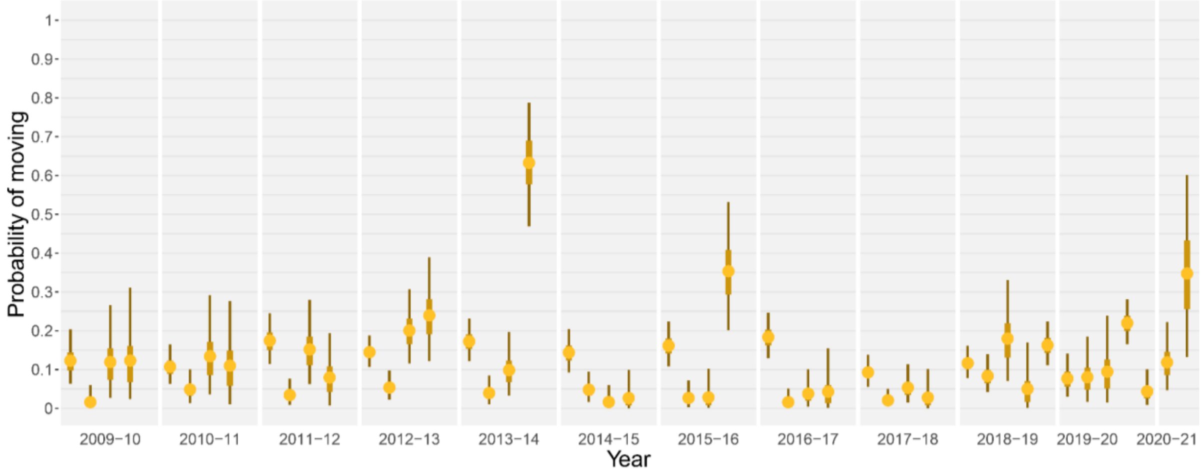


**Figure S11**. Probability of moving to migratory area 3 (yellow area on fig. S1), conditional on departure. This probability is constant across sexes. In each year, the four successive points correspond to the four successive winter occasions. Point estimates are posterior means, inner and outer line segments indicate 50% and 95% credible intervals.

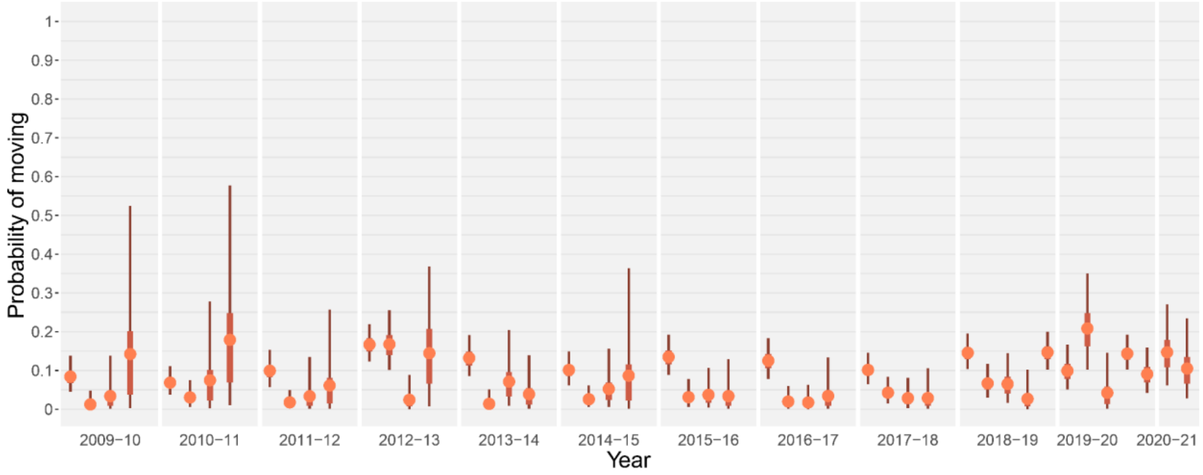


**Figure S12**. Probability of moving to migratory area 4 (pink area on fig. S1), conditional on departure. This probability is constant across sexes. In each year, the four successive points correspond to the four successive winter occasions. Point estimates are posterior means, inner and outer line segments indicate 50% and 95% credible intervals.

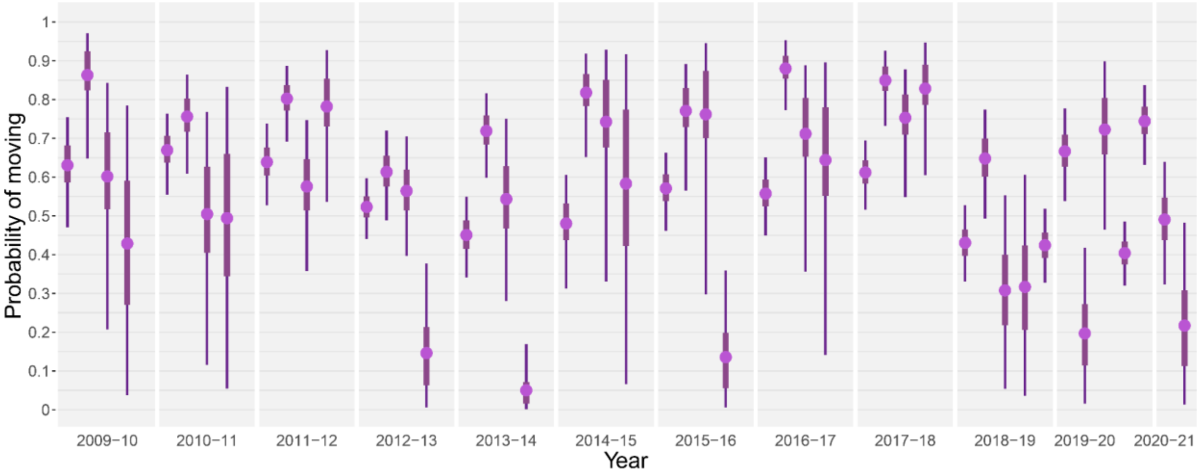


**Figure S13**. Probability of moving to migratory area 5 (the 'ghost area'), conditional on departure. This probability is constant across sexes. In each year, the four successive points correspond to the four successive winter occasions. Point estimates are posterior means, inner and outer line segments indicate 50% and 95% credible intervals.

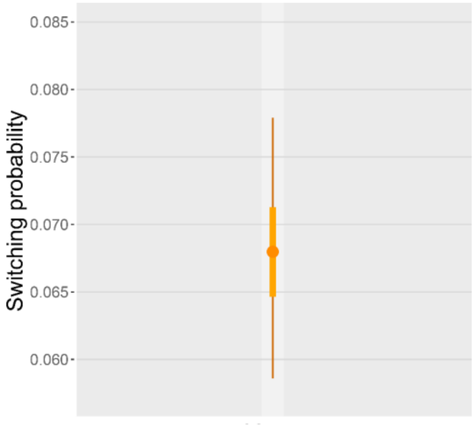


**Figure S14**. Probability of switching to another migratory area (i.e. not remaining in the same migratory area). This probability is constant across occasions, years, and sexes. Points are posterior means, inner and outer line segments indicate 50% and 95% credible intervals.

*Detection probabilities*

Hereafter we provide graphical summaries of the posterior distribution of the resighting probability in the residency area (fig. S15) and observed migratory areas (fig. S16 to S19).

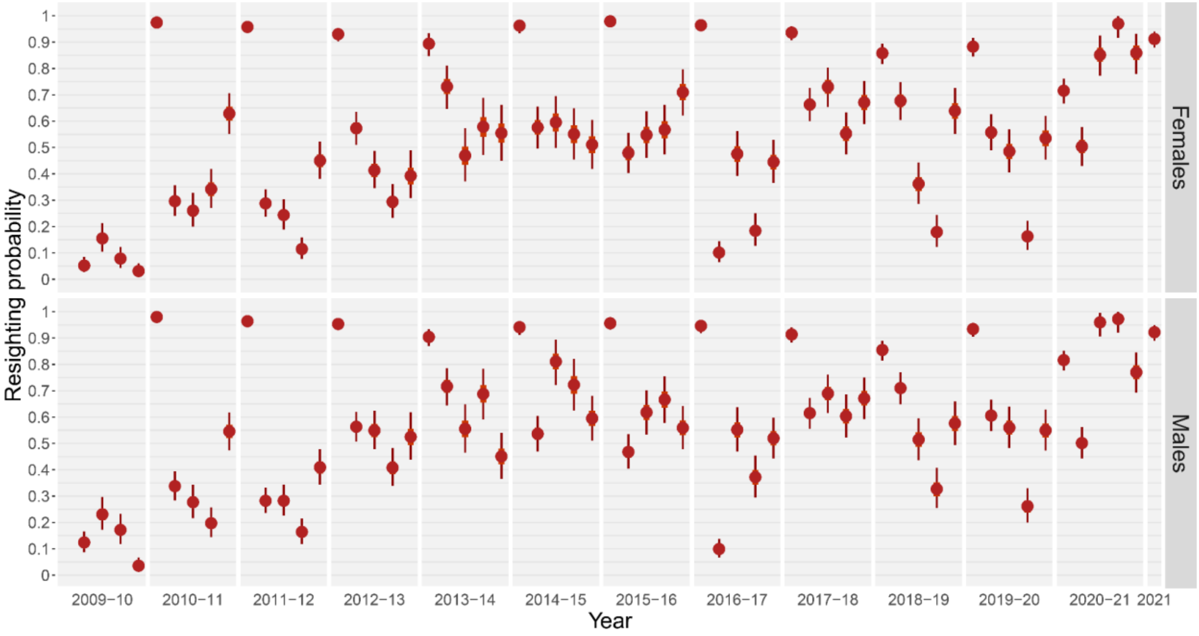


**Figure S15**. Resighting probability in the residency area in females and males. Point estimates are posterior means, inner and outer line segments indicate 50% and 95% credible intervals. In each year, the five successive points correspond to the five successive occasions: the breeding season and the four winter occasions. The very first point (breeding season, year 1) is not shown because all present individuals are entering the dataset and hence sighted with certainty.

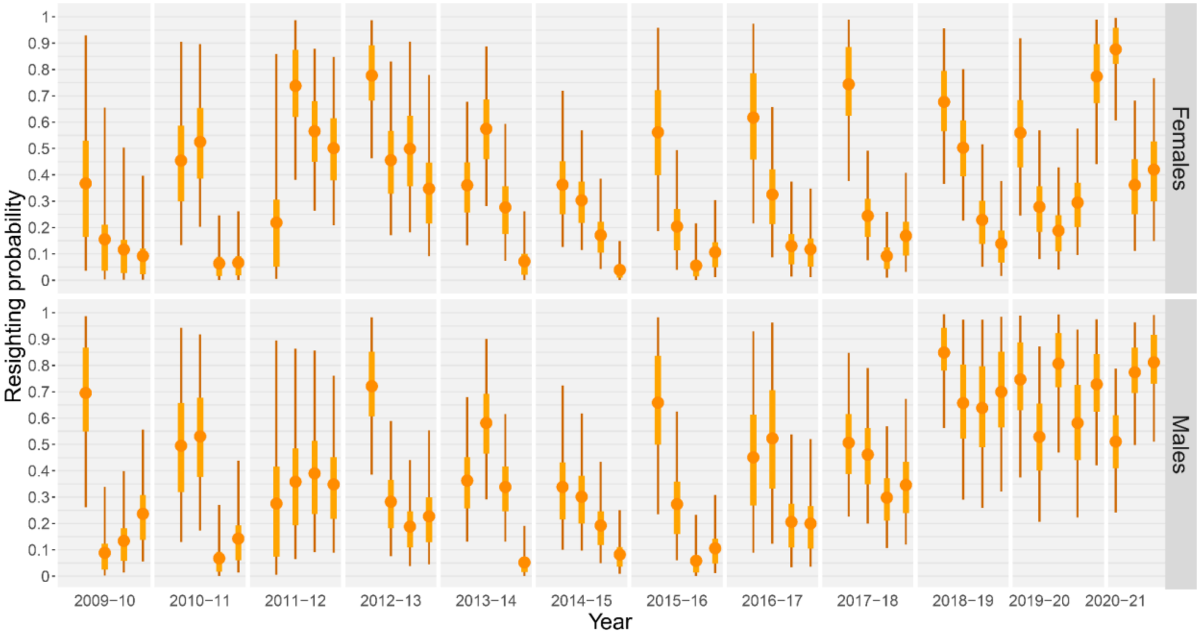


**Figure S16**. Resighting probability in migratory area 1 (orange points on fig. S1) in females and males. In each year, the four successive points correspond to the four successive winter occasions. Point estimates are posterior means, inner and outer line segments indicate 50% and 95% credible intervals.

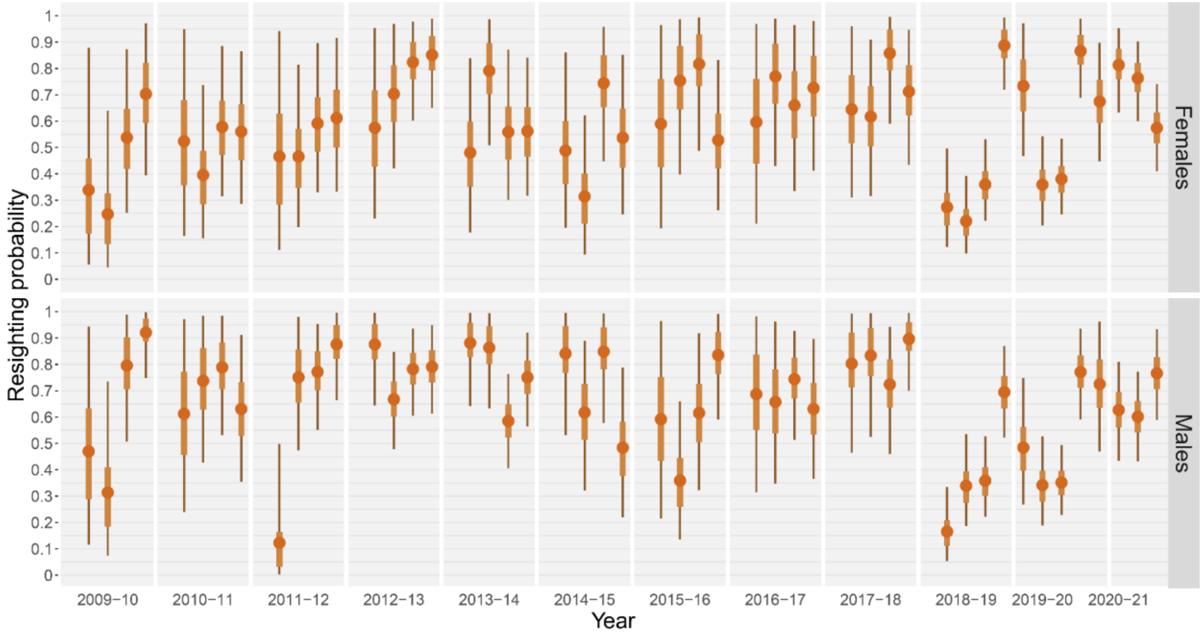


**Figure S17**. Resighting probability in migratory area 2 (brown area on fig. S1) in females and males. In each year, the four successive points correspond to the four successive winter occasions. Point estimates are posterior means, inner and outer line segments indicate 50% and 95% credible intervals.

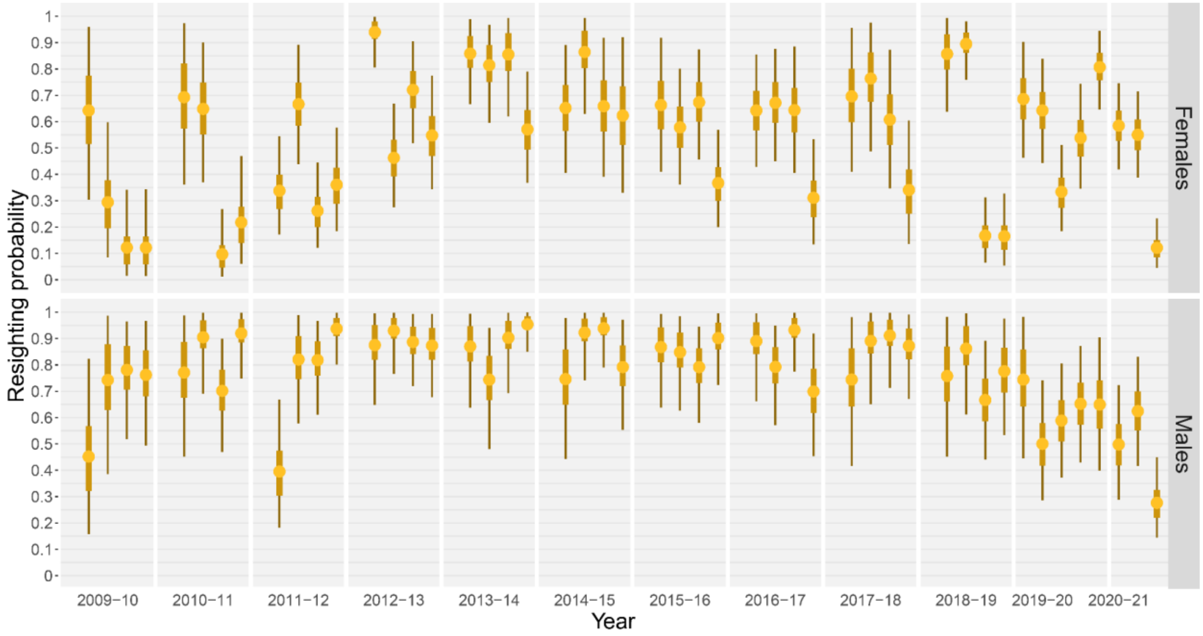


**Figure S18**. Resighting probability in migratory area 3 (yellow area on fig. S1) in females and males. In each year, the four successive points correspond to the four successive winter occasions. Point estimates are posterior means, inner and outer line segments indicate 50% and 95% credible intervals.

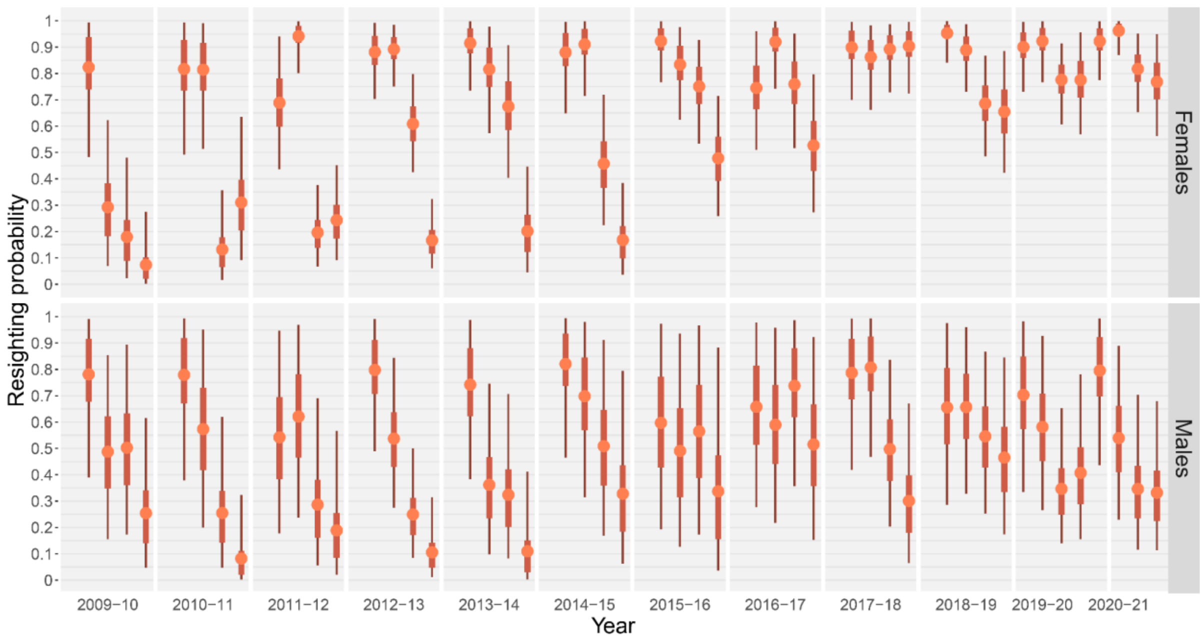


**Figure S19**. Resighting probability in migratory area 4 (pink area on fig. S1) in females and males. In each year, the four successive points correspond to the four successive winter occasions. Point estimates are posterior means, inner and outer line segments indicate 50% and 95% credible intervals.

### Part S5
